# HeartMap: An Integrated Cell Atlas of 2.4 million cells across 209 Individuals in Health and Disease

**DOI:** 10.1101/2025.11.06.686995

**Authors:** Yesh Datar, Mark Chaffin, Bridget Simonson, Alex Mandia, Patrick T. Ellinor

**Affiliations:** Cardiovascular Disease Initiative, The Broad Institute of MIT and Harvard, Cambridge, MA, USA; Boston University Chobanian and Avedesian School of Medicine, Boston, MA USA; Cardiology Division, Mass General Brigham, Boston, MA, USA; Heart and Vascular Institute, Mass General Brigham, Boston, MA, USA

## Abstract

Cardiovascular disease remains the leading cause of global mortality. Understanding its complexity requires dissecting the heart's cellular landscape. We present HeartMap, a comprehensive single-nucleus RNA sequencing atlas of the adult human heart. This resource integrates data from nine studies, encompassing over 2.4 million nuclei, 209 individuals, eight anatomical regions, and seven disease and healthy states. After rigorous data harmonization and benchmarking of batch-correction methods, we characterized transcriptional diversity across 14 cell types. To demonstrate the utility of HeartMap, we identified robust disease-associated gene signatures in dilated cardiomyopathy by comparing across multiple studies. Notably, we identified distinct activated fibroblast populations, with COL22A1 or TNC enrichment, revealing potential differences between inherited and ischemic cardiomyopathies. HeartMap provides a valuable tool for exploring cardiac disease at the single-cell level, facilitating both fundamental research and potential therapeutic development.

## Main

Recent advances in single-cell and single-nucleus RNA sequencing (snRNA-seq) technologies have enabled extensive transcriptomic analyses of human organ-tissue systems across a variety of diseases.^1–4^ These analyses have led to an unparalleled study of diseases by looking at the changes in common and rare cell populations. Atlases allow researchers to interrogate cell populations at scale to identify disease-associated genes, cell-to-cell interactions, and biomarkers to detect disease. Integrated cell atlases, which combine multiple RNA-seq studies together, allow for even more robust comparisons between studies and increase the statistical power for downstream analyses.^4^

To date, many research groups have studied cardiovascular disease in healthy and disease tissues.^5–12^ While some variation in gene expression and cell states can be explained by the specific disease under study, substantial variability is introduced based on sequencing protocols, data pre-processing, and downstream analysis methods. A well-integrated heart disease cell atlas would help reduce technical noise and augment meaningful biological signals from these studies. Furthermore, an atlas of this size would allow for unprecedented transcriptomic comparisons between cardiomyopathies, regional cardiac anatomies, cell types, and biological sex.

In the current work, we constructed an integrated atlas through the re-processing of published studies and robust batch-correction using previously described packages for benchmarking and integration. Furthermore, we conducted pairwise differential expression tests between different disease states, while controlling for study-specific effects. To identify a robust set of genes associated with dilated cardiomyopathy, we performed differential expression tests comparing dilated cardiomyopathy versus control hearts across three independent studies. We discovered disease-associated genes expressed in various cell types that are consistent across studies. Finally, we investigated specific cell populations and identified two activated fibroblast niches that are enriched in primary and ischemic cardiomyopathies.

## Results

### Published datasets form the foundation of HeartMap

We collected published datasets from 9 snRNA-seq studies to create HeartMap, including 8 published and 1 unpublished works. ^5–12^ Combined, these datasets included 485 healthy and diseased heart samples from various anatomies of 209 individuals (**Table 1**).

**Table 1.**
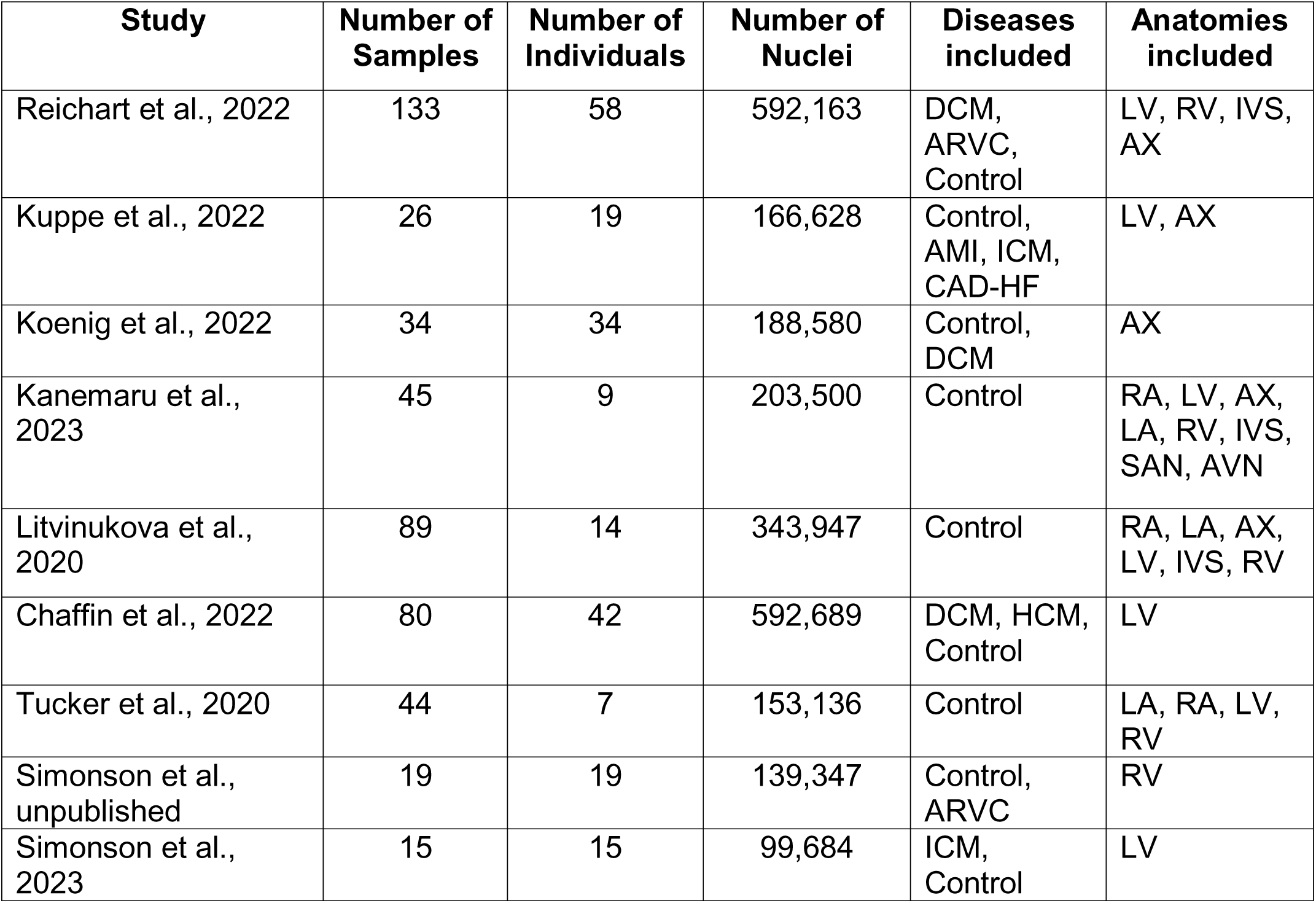
Number of samples, individuals, post-quality control nuclei, disease phenotypes, and anatomical regions included in each study. Note that multiple individuals have technical replicates in each study. The following acronyms are used here: healthy (Control), hypertrophic cardiomyopathy (HCM), dilated cardiomyopathy (DCM), arrhythmogenic cardiomyopathy (ARVC), coronary artery disease (CAD), acute myocardial infarction (AMI), ischemic cardiomyopathy (ICM), left ventricle (LV), left atrium (LA), left ventricular apex (AX), right ventricle (RV), right atrium (RV), interventricular septum (IVS), sinoatrial node (SAN), and atrioventricular node (AVN).

| Study | Number of Samples | Number of Individuals | Number of Nuclei | Diseases included | Anatomies included |
| --- | --- | --- | --- | --- | --- |
| Reichart et al., 2022 | 133 | 58 | 592,163 | DCM, ARVC, Control | LV, RV, IVS, AX |
| Kuppe et al., 2022 | 26 | 19 | 166,628 | Control, AMI, ICM, CAD-HF | LV, AX |
| Koenig et al., 2022 | 34 | 34 | 188,580 | Control, DCM | AX |
| Kanemaru et al., 2023 | 45 | 9 | 203,500 | Control | RA, LV, AX, LA, RV, IVS, SAN, AVN |
| Litvinukova et al., 2020 | 89 | 14 | 343,947 | Control | RA, LA, AX, LV, IVS, RV |
| Chaffin et al., 2022 | 80 | 42 | 592,689 | DCM, HCM, Control | LV |
| Tucker et al., 2020 | 44 | 7 | 153,136 | Control | LA, RA, LV, RV |
| Simonson et al., unpublished | 19 | 19 | 139,347 | Control, ARVC | RV |
| Simonson et al., 2023 | 15 | 15 | 99,684 | ICM, Control | LV |

We included over 2.4 million nuclei from 7 disease and non-disease states including dilated cardiomyopathy (DCM), hypertrophic cardiomyopathy (HCM), ischemic cardiomyopathy (ICM), arrhythmogenic right ventricular cardiomyopathy (ARVC), acute myocardial infarction (AMI), coronary artery disease (CAD-HF), and non-failing control hearts (Control). The dataset includes 8 cardiac anatomies including the left and right atria and ventricles (LV, LA, RV, RA), left ventricular apex (AX), interventricular septum (IVS), sinoatrial (SA), and atrioventricular (AV) nodes.

Not all clinical metadata was made available, however of those provided, the individuals had a median age of 54 years (IQR: 45-62, n = 208 individuals), 39% were female. Apart from the non-failing and acutely infarcted, all samples from studies were from hearts in chronic, end stage disease. This atlas provides a new resource for comparing multiple diseases to controls and serves as a transcriptomic reference for studying cardiovascular disease (**Figure 1**).

**Figure 1.**
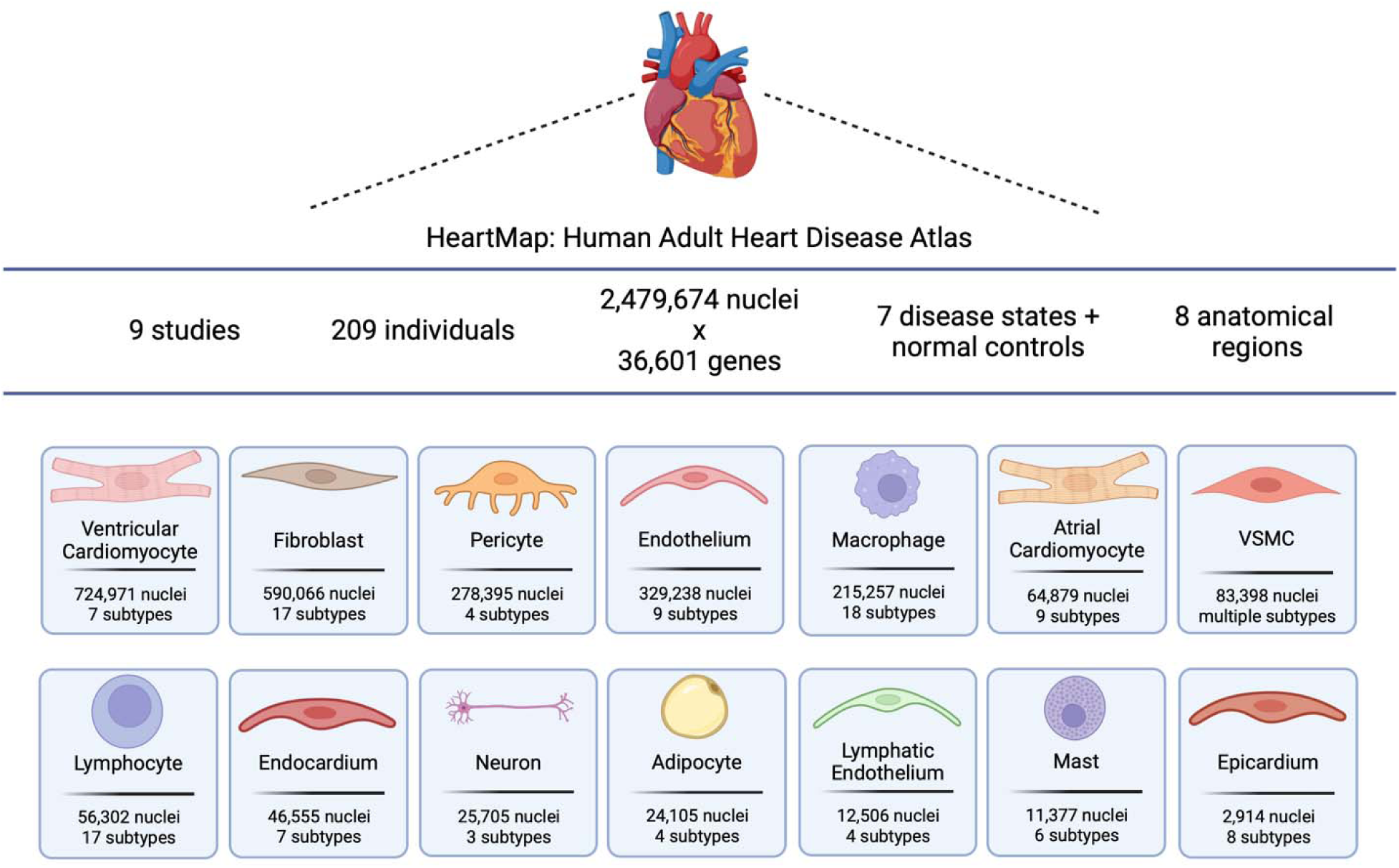
Schematic overview of the composition of HeartMap. The number of nuclei and sub-clusters assigned to each cell type based on a global map is provided (Created with Biorender.com).

### Data integration across studies controls for batch-effects in HeartMap

We removed sample duplicates and incorrectly labelled samples from each of the studies prior to any batch-correction and downstream analysis (**see Methods**). For our batch-correction across studies, we considered up to 4 parameterizations of the following integration methods: 1. Harmony^13^, 2. Scanorama^14^, 3. single-cell Variational Inference (scVI)^15^, and (4) single-cell ANnotation using Variational Inference (scANVI)^16^. We benchmarked the performance of each approach using Single-cell integration benchmarking (scIB).^17^ We considered integration performance across both study and individual, separately. Notably, individuals represent distinct patients and may have multiple samples if multiple tissues were surveyed, or they were included in multiple studies. scANVI and Harmony revealed the highest scores for biological conservation and batch correction, respectively (**Supplemental Figure 1**); however, given that scANVI had the overall best performance when considering the preservation of biological signal while correcting for batch effects, this method was chosen to construct the complete atlas.

After harmonizing the atlas with scANVI, we visually assessed nuclei mixing by individual of origin and study (**Figure 2**). Large transcriptional variation has been shown to exist between individuals, even within cell types, most notably in cardiomyocytes, as seen in Tucker et al. (Supplemental Figure 2).^7^ This can be seen in the variation that exists in the uniform manifold approximation and projection (UMAP) representation of the uncorrected atlas by individual and study (**Figure 2A,C**) compared to the corrected atlas by individual and study (**Figure 2D,F**). After correction, cell types resolve in a harmonized fashion reflecting appropriate batch-correction on a subjective level with objective metrics provided in the benchmarking metrics.

**Figure 2.**
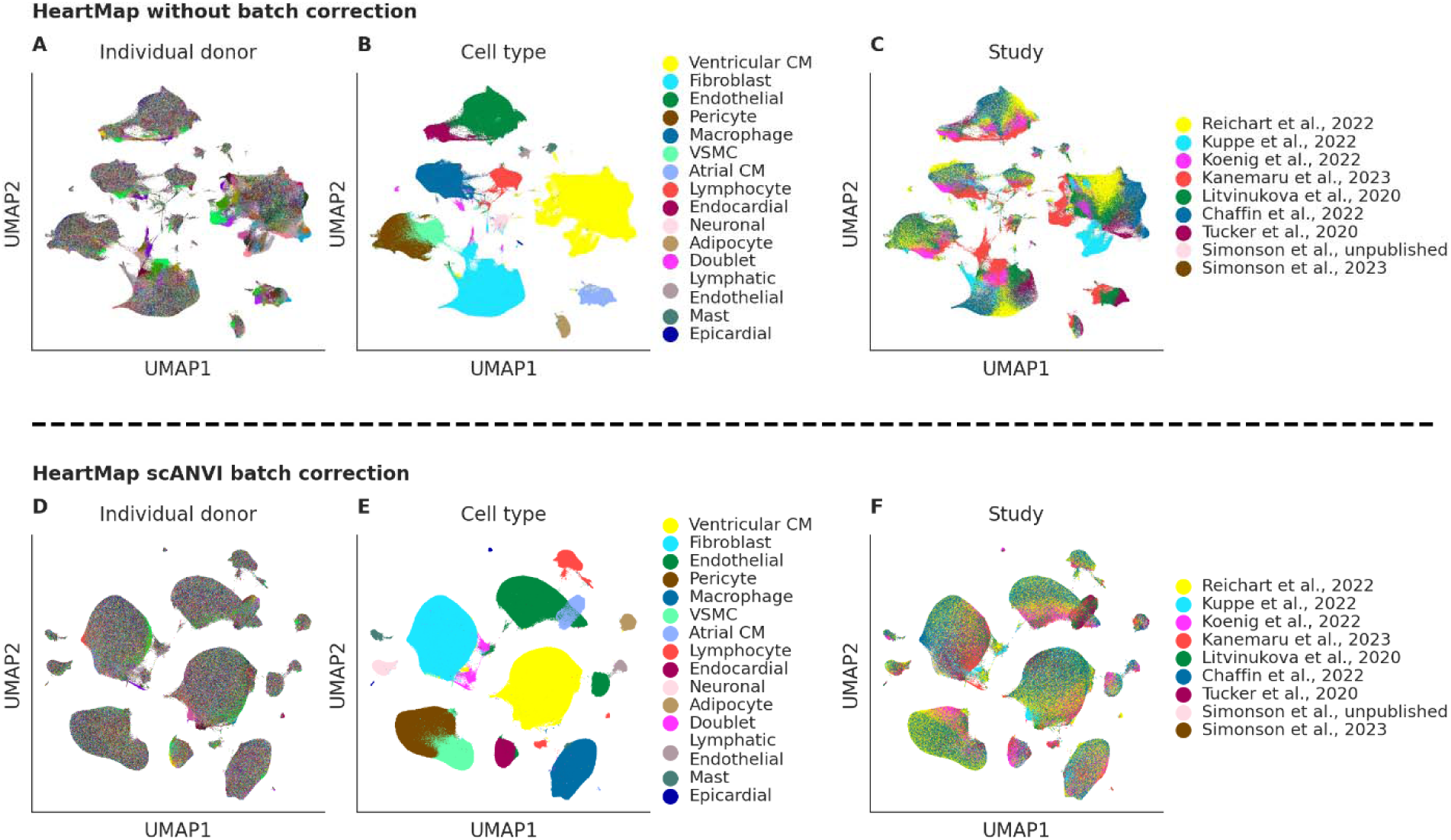
Batch-correction on the HeartMap atlas using scANVI. **A,B,C**, A Uniform Manifold Approximation and Projection (UMAP) representation of all nuclei without batch correction colored by individual (**A**), cell type (**B**), and study (**C**). **D,E,F,** A UMAP representation of all nuclei with batch correction from scANVI colored by individual (**D**), cell type (**E**), and study (**F**). CM, cardiomyocyte; VSMC, vascular smooth muscle cell.

### Mapping of the cardiac cell landscape reveals compositional shifts in common and rare cell types across disease

After data integration and quality control, we mapped the combined 2,479,674 nuclei with scANVI (see **Methods**). The final atlas revealed 52 clusters representing 14 known common and rare cell types in heart tissue. Additional distinctions based on both anatomical region and disease were also observed (**Supplemental Figure 2A**). We identified distinct marker genes for all cell types, many of which are known canonical markers as expected (**Supplemental Figure 2B, Supplemental Table 1)**.

After restricting to anatomical regions captured by at least one disease state, we observed differences in global cell type proportions by disease. For example, cardiomyocyte proportions have a credible increase in healthy (e.g., NF control) versus disease and chronic states (e.g., DCM, HCM, ARVC, ICM, CAD-HF), which recapitulate known findings (**Figure 3**).^18^ Additionally, we considered whether any cell type proportions varied markedly by study. Reassuringly, proportions appeared generally similar, or notable differences could be explained by the fraction of non-failing control samples in each study or the inclusion of different anatomical regions. For example, Tucker et al., 2020, Kanemaru et al., 2023 and Litvinukova et al., 2023 are the only studies that included atrial tissue, and reassuringly atrial cardiomyocytes are only found in these studies (**Supplemental Figure 3)**.

**Figure 3.**
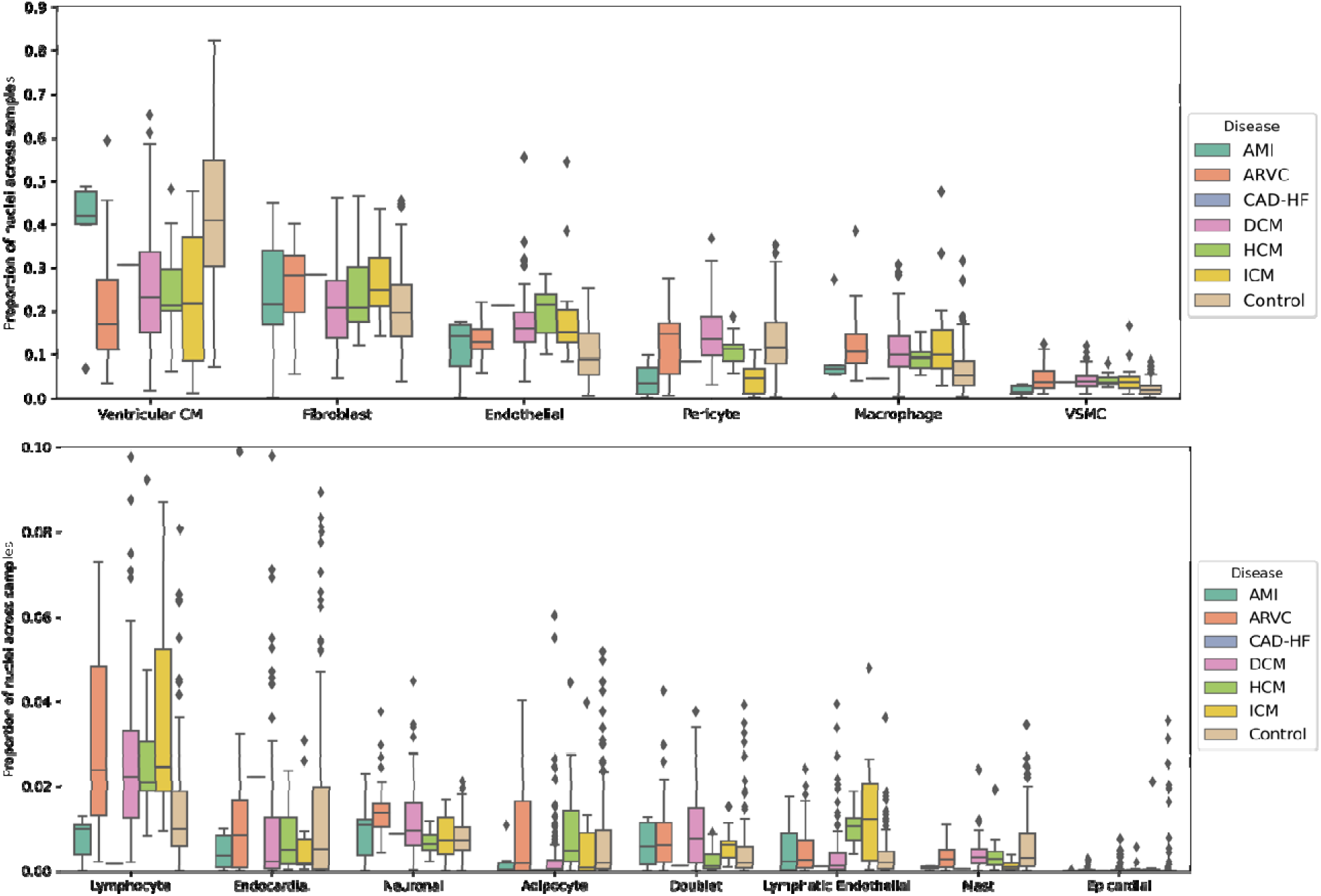
Cell type composition by disease. Box and whisker plots show the distribution of cell type proportions across all samples, with the center of the box being the median and lower and upper bounds being the 25^th^ and 75^th^ percentiles, respectively. Outliers are shown as points in the box plots. Sample is considered as a unique combination of “study” + “individual” + “anatomy” (see Methods). Anatomies only present in one disease state including atrioventricular node, sinoatrial node, right atrial, and left atrial cells have been removed. DCM, dilated cardiomyopathy; HCM, hypertrophic cardiomyopathy; ARVC, arrhythmogenic right ventricular cardiomyopathy; ICM, ischemic cardiomyopathy; AMI, acute myocardial infarction; CM, cardiomyocyte; VSMC, vascular smooth muscle cell.

### Transcriptional changes distinguish the diseased and healthy heart

To explore inter-sample heterogeneity in gene expression across all cell types, we constructed a sample-level UMAP embedding by summing the expression of all nuclei from that sample (i.e., unique combination of individual, study, and chamber). When visualizing this global atlas UMAP representation based on study, anatomy, sex-assignment, and disease phenotype, we noted that samples separated by phenotypic characteristics. For example, samples strongly separated by sex based on the first two UMAP dimensions. Additionally, atrial, SA and, AV node samples separated from the ventricular and IVS samples (**Figure 4B**). Notably, control samples separate from diseased samples (**Figure 4D**). AMI samples from individuals P2, P3, P6, and P9 collected from days of infarct between 2-5 days were closer to control samples than P10 and P15 which were collected at days 6 and 11, respectively.

**Figure 4.**
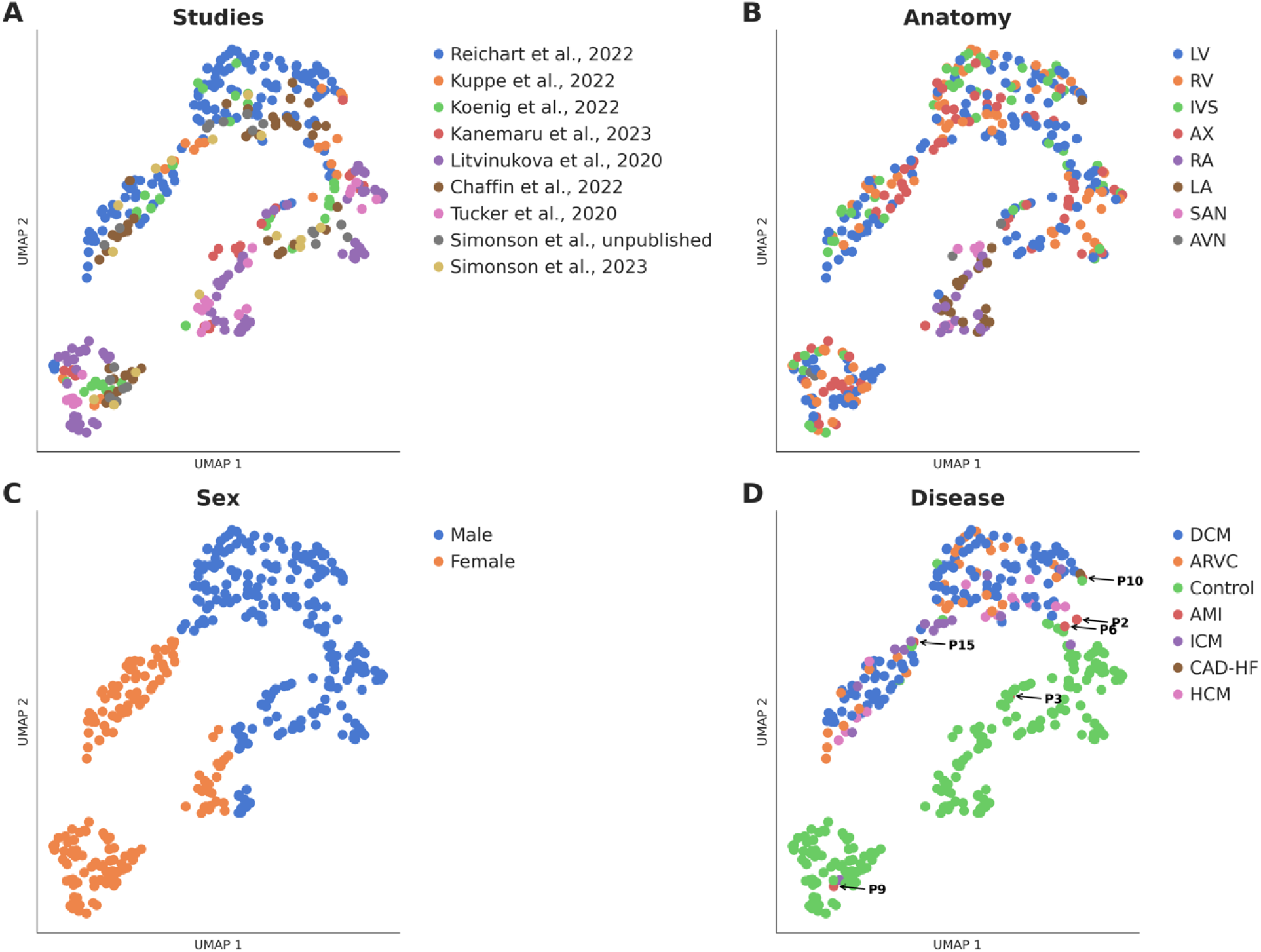
Sample-level embeddings of HeartMap. **A, B, C, D,** A UMAP representations of the global atlas with annotations based on study of origin (**A**), anatomy (**B**), sex-assignments (**C**), and disease (**D**). Each point in the UMAP represents one sample in the study. In plot **D**), samples with acute myocardial infarctions are labelled. LV, left ventricle; RV, right ventricle; LA, left atrium; RA, right atrium; AX, left ventricular apex; IVS, interventricular septum; SAN, sinoatrial node; AVN, atrioventricular node; DCM, dilated cardiomyopathy; HCM, hypertrophic cardiomyopathy; ARVC, arrhythmogenic right ventricular cardiomyopathy; ICM, ischemic cardiomyopathy; AMI, acute myocardial infarction.

We also constructed sample-level UMAP embeddings of each cell type level to distinguish transcriptional differences between sex-assignment, disease, study, and anatomy. Sex-assignment primarily drove sample-level differences (**Supplemental Figure 4A)**. When examining the first two UMAP dimensions, control samples separated from diseased samples in common cell types while rarer cell types had a less clear distinction (**Supplemental Figure 4B**).

In conjunction with the findings on the global and cell-level embedding maps, we ran differential expression tests to investigate the transcriptomic differences between diseases and sex-assignments. Using pairwise comparisons between disease groups, we found that the most differential genes were in common cell types such as ventricular cardiomyocytes and fibroblasts **(Figure 5)**. Additionally, we observed more differentially expressed genes in diseased versus control comparisons than disease versus disease comparisons. Notably, we still saw an appreciable, although relatively fewer, number of differentially expressed genes between inherited cardiomyopathy (DCM vs HCM) and ischemic cardiomyopathy (ICM vs AMI) across cell types as well. In the pairwise comparison between female and male sex-assignment groups, we found that most genes with large effect estimates were Y or X chromosome specific and the highest number of these genes were in ventricular cardiomyocytes and fibroblasts **(Supplemental Table 2).**

**Figure 5.**
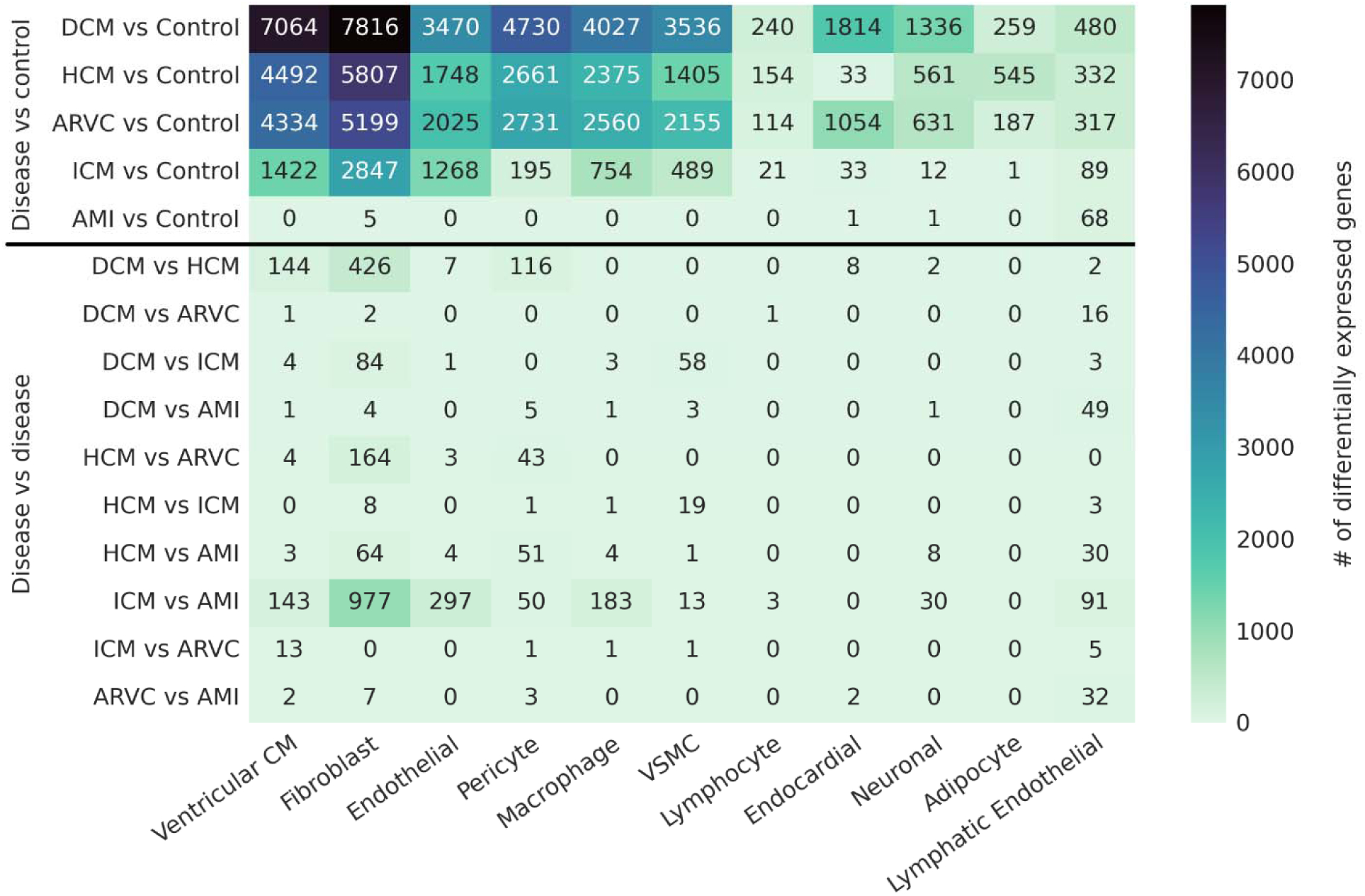
Heatmap of the differentially expressed genes in each pairwise phenotypic comparison. For each disease versus control or disease versus disease comparison, the number of differentially expressed genes in each cell type are shown. Differentially expressed genes are defined as having FDR-adjusted P < 0.05 based on both CellBender and CellRanger counts, a low chance of being from background contamination (background heuristic < 0.5), and a concordant direction of effect in both CellBender and CellRanger. DCM, dilated cardiomyopathy; HCM, hypertrophic cardiomyopathy; ARVC, arrhythmogenic right ventricular cardiomyopathy; ICM, ischemic cardiomyopathy; AMI, acute myocardial infarction; CM, cardiomyocyte; VSMC, vascular smooth muscle cell.

### Disease-associated genes in dilated cardiomyopathy are replicated between studies

HeartMap also enables the comparison of differential expression (DE) tests between published studies. Examining disease-associated genes across multiple studies allows us to focus on the most prominent disease-associated signals and decreases the background signals generated from individual studies alone. In this analysis, we focused on the comparison of DCM to NF control hearts as DCM was captured in three distinct studies and allowed for a robust comparison of effect estimates. We performed DE tests for the three reprocessed studies in our atlas (Chaffin et al., 2022, Reichart et al., 2022, and Koenig et al., 2022) which contained end-stage DCM and NF control samples (see **Methods**).

As expected, we saw a strong concordant relationship when comparing the within study effect estimates between the DE test from published data and our reprocessed data **(Supplemental Figure 5A,B)**. This gave reassurance that we had not significantly changed the underlying data during reprocessing. Across the three DE tests from the reprocessed studies, we identified a set of genes with a strong positive or negative logFC estimate (|logFC| > 1.5) across multiple cell types, primarily in ventricular cardiomyocytes, fibroblasts, and endocardial cells (**Figure 6, Supplemental Table 3)**.

**Figure 6.**
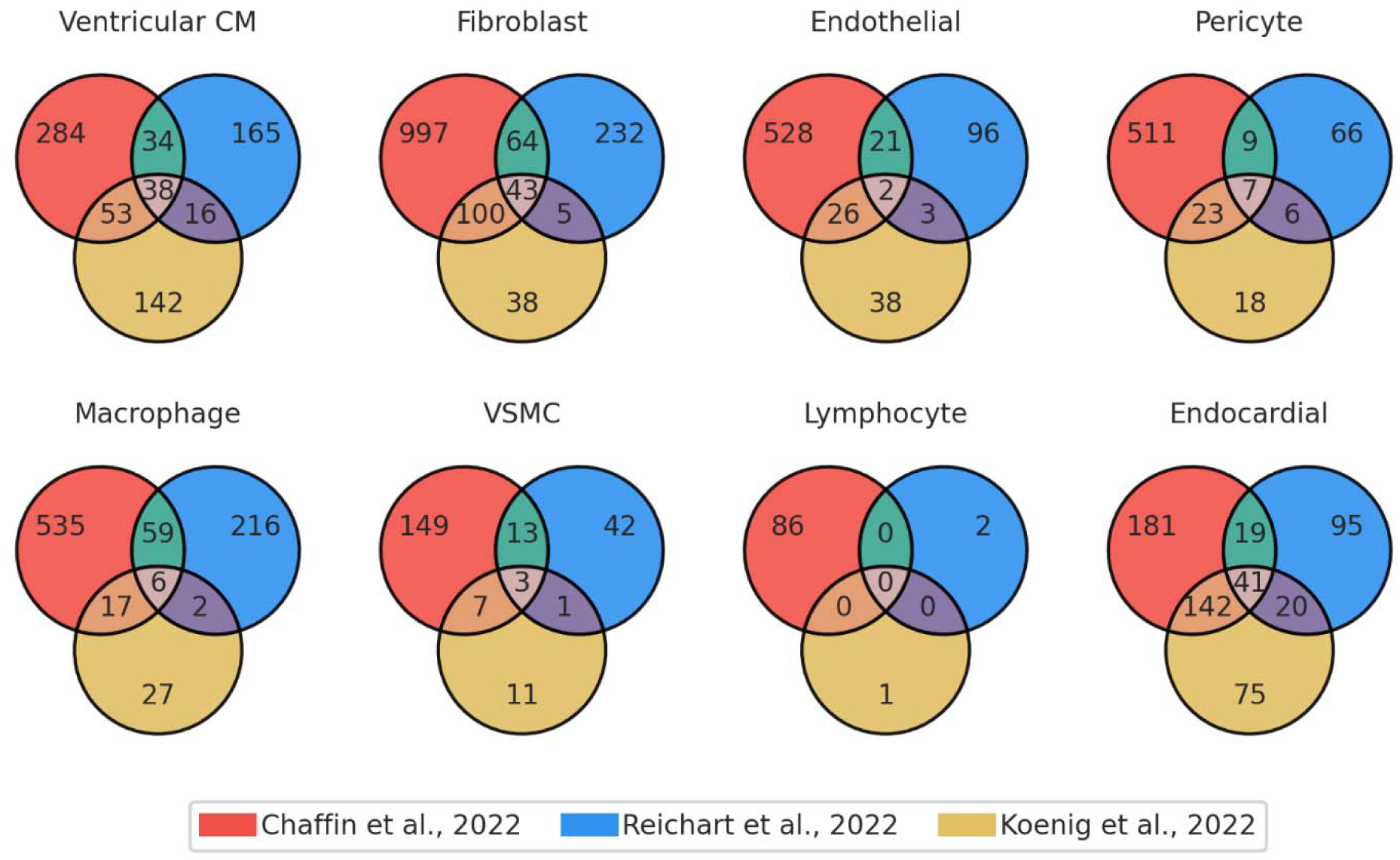
Venn diagrams demonstrating the overlap of strongly differentially expressed genes in DCM vs NF control samples across multiple re-processed studies. For each cell type, strongly differentially expressed genes (|logFC| > 1.5, FDR-adjusted P-value < 0.05, and background heuristic < 0.5 in both comparator groups) were identified in each study separately. Genes with directionally consistent effects between studies are shown in the overlap. In cases where a gene had a strong, but directionally inconsistent effect between studies, it was included in the tabulation for each study-specific count but excluded from the intersection. CM, cardiomyocyte; VSMC, vascular smooth muscle cell.

In ventricular cardiomyocytes, some example genes with strong upregulation in DCM across all three studies included: *UNC80, PLCE1, NPPA,* and *NPPB*. Interestingly, *UNC80* mutation has been associated with a syndrome of hypotonia, dyskinesia, and dysmorphism.^19^ Similarly, *PLCE1* plays roles in modulating beta-adrenergic receptor dependent cardiac contraction and inhibits hypertrophy.^20^ Additionally, *NPPA* and *NPPB* encode atrial and B-type natriuretic peptides and are classical biomarkers of heart failure.^21^ Strongly downregulated genes in DCM across all three studies included *BMP7* which has been shown to be activated by *YY1* and co-modulate *CTGF* to suppress *LMNA* DCM and cardiac fibrosis.^22^

In endothelial cells, *AC093817* was strongly upregulated in DCM across all studies and has been implicated as a potential biomarker of DCM from plasma samples.^23^ Strongly upregulated genes in endocardial cells included *RND3, BMP4/6,* and *ELN*. *Rnd3*(+/−)) mice have been shown to be predisposed to developing dilated cardiomyopathy with heart failure after transverse aortic constriction stress compared to normal mice.^24^ Bone morphogenetic proteins (*BMP*) including *BMP4/6* have also been implicated in DCM by influencing cardiac development.^25,26^ Elastin (*ELN*) has been shown to have higher elastin-collagen ratio in the endocardium and is crucial for recoil during the systolic contraction.^27^

Finally, in pericytes several genes showed strong downregulation in DCM across studies. This included *ADAMTS4* which has been shown, through inhibition, to prevent cardiac fibrosis by reducing the release of TGF-β, a transforming growth factor, due to EDA-fibronectin cleavage.^28^ Additional cell types and associated genes are presented in **Supplemental Table 3.** Pathway analysis revealed that there were multiple common pathways between the three studies primarily in fibroblasts, pericytes, and macrophages (**Supplemental Figure 6, Supplemental Table 4**).

Despite an overall positive correlation between gene effect estimates in the comparison of DCM to control between studies, there exists substantial heterogeneity (**Supplemental Figure 5C**). To assess the degree of heterogeneity across studies, we calculated an I^2^ coefficient for each gene tested in each cell type. A high percentage of genes tested had an I^2^ >75% in each cell type (roughly 40% ventricular cardiomyocytes, 50% fibroblasts, 30% pericytes, 37% endothelial, 37% macrophage, 28% lymphocyte, 18% endocardial), indicating high heterogeneity. While the above analysis focused on genes with consistent effects across studies, it is unclear why a large fraction of genes shows very disparate effects between DCM and control between these studies. Genes with higher I^2^ coefficients did not seem to be associated with overall expression levels, the degree to which the gene is likely driven by background RNA, or the enrichment of specific and unique pathways.

### Distinct populations of activated fibroblasts in inherited and ischemic cardiomyopathies

HeartMap allows for investigation into cell types after rigorous batch-correction to account for individual- and study-level differences. We systemically sub-clustered all major cell types and identified between 3 and 18 sub-clusters across cell types (**Supplemental Figures 7**-**20**). Of particular interest, we identified 17 sub-clusters of fibroblasts across the 590,066 fibroblast nuclei in HeartMap (**Figure 7A**). Three of these sub-clusters, 0, 7 and 9, were characterized as activated fibroblast populations based on elevated expression of canonical genes, including *POSTN* and *FAP*. However, populations 7 and 9 diverged in their expression of additional markers such as *COL22A1* and *TNC*, respectively, suggesting these may be distinct fibroblast populations (**Figure 7B,C**).

**Figure 7.**
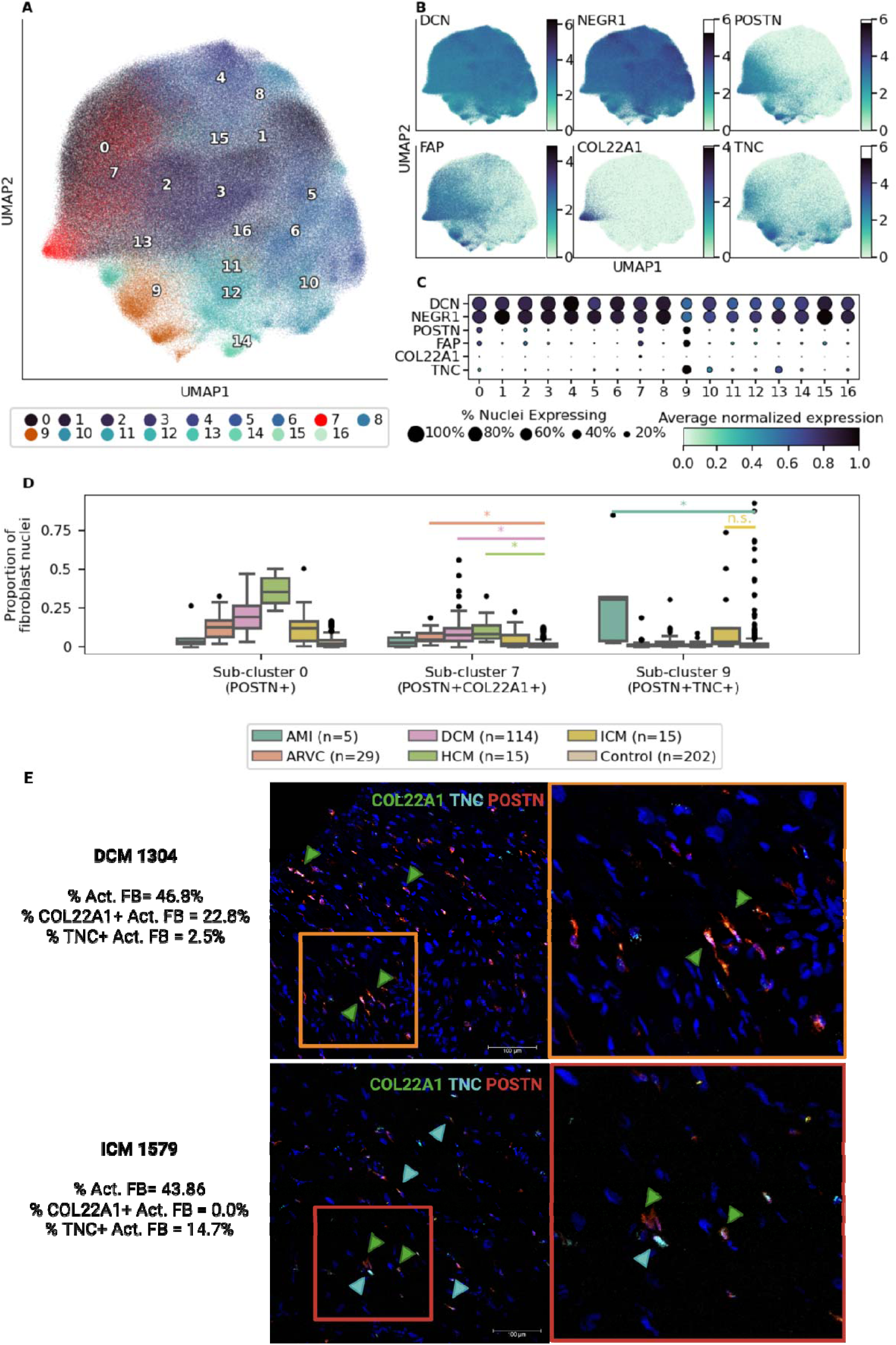
Fibroblast investigation based on sub-clustering and compositional analysis. **A**, UMAP plot of 590,066 fibroblast nuclei colored by sub-cluster. The *COL22A1*-enriched sub-cluster is colored in red (cluster 7) and the *TNC*-enriched sub-cluster is colored in dark orange (cluster 9). **B**, UMAP with overlay of log-normalized expression for canonical fibroblast genes (*DCN*, *NEGR1*), general activated fibroblast genes (*POSTN, FAP)*, and sub-cluster specific genes (*COL22A1*, *TNC*). **C,** Dotplot depicting the percent of nuclei in each sub-cluster expressing the genes in **B** (size) and the average log-normalized expression, normalized by gene across sub-cluster (shade). **D**, Box plots where samples are plotted by the proportion of fibroblast nuclei expressed in their respective sub-clusters. Samples are identified by a unique combination of individual-study-anatomy (see Methods) and number of samples per disease are listed in the legend. “*” denotes credible changes between control and diseased samples. “n.s.” denotes no significant credible change was noted. **E**, Representative *in situ* hybridization Image validation from two LV tissue samples. The computationally predicted percentage of activated fibroblasts in each sample are included with labels of these cell populations in the images. Markers are for activated fibroblasts (*POSTN*, red), *TNC* (aqua) and *COL22A1* (green). Green arrows represent POSTN+COL22A1+ cells, aqua arrows represent POSTN+TNC+ cells. Scale bar = 100 um.

Of those activated fibroblast populations, compositional analysis revealed that sub-clusters 7 and 9 were more prevalent in inherited/primary cardiomyopathies (i.e. DCM, HCM, ARVC) and ischemic disease (i.e. AMI, CAD-HF, and ICM), respectively (**Figure 7D**). Genes upregulated in these two activated fibroblast populations show gene set enrichment for collagen fibril organization (**Supplemental Table 5)**. Additionally, activated fibroblast populations in the inherited cardiomyopathy group (sub-cluster 7) show gene enrichment for bone development, skeletal system development and cardiac hypertrophy with marker genes such as *POSTN*, *FAP*, and *COL22A1*.

Activated fibroblasts in the ischemic disease group (sub-cluster 9) show gene enrichment for granulocyte chemotaxis, neutrophil migration, and wound healing with marker genes such as *POSTN*, *FAP*, and *TNC*. To experimentally validate these activated fibroblast populations, we carried out RNAscope for *COL22A1* (cluster 7) and *TNC* (cluster 9), using *POSTN* as a marker for activated fibroblasts. We selected 3 NF, 3 ICM and 3 inherited cardiomyopathy samples by specific criteria (see **Methods**) and confirmed that these were two separate cell populations and that the proportions of each cell population in each sample were similar between computationally predictions and experimental findings (**Figure 7E**, **Supplemental Table 6**).

## Discussion

In the present study, we have constructed HeartMap, the largest single-nucleus atlas used to study human adult heart disease. This map allows us to characterize common and rare cell types across various disease and healthy phenotypes. We demonstrate that control and diseased human samples separate based on transcriptomic profiles based on disease, anatomy, and sex-assignment. Furthermore, our pairwise DE test between diseases and control samples reveal more genes separating disease from control than disease from other diseases. Our analysis of disease-associated genes across studies in DCM shows that there are many common up- and down-regulated genes and pathways in multiple cell types; however, despite sharing some similarities, effect estimates were often discordant across studies. Finally, we highlight the usefulness of our atlas to look at cell types at a larger scale than previously possible with disease-based comparisons. We show that fibroblasts have unique transcriptional differences in inherited and ischemic cardiomyopathies with markers associated with activated, collagen-depositing subtypes.

After careful re-processing and batch-correction, we showed that we can robustly harmonize data across multiple studies to maintain meaningful biological signals in the highly heterogenous tissue of the heart. We found genes that serve as strong markers for each cell type across multiple studies, allowing us to better characterize a global transcriptional signature of the heart in healthy and disease states. Within these cell types, genes show strong dysregulation in disease states compared to controls; however, there exists a smaller subset of genes differentiating disease states from one another. These between-disease comparisons are only possible with the aggregation of multiple studies as performed in the construction of HeartMap. Differentiating the transcriptional signatures between cardiovascular disease groups may aid in precision medicine approaches by informing further work on biomarker and therapeutic target discovery.

In the same vein, we also present an analysis of disease-associated genes in DCM across three independent studies. Combining samples from multiple studies improves statistical power and allows for the identification of additional DCM genes. Additionally, we also showed that effect estimates are not always consistent between studies. This demonstrates that: 1) batch effects are critical to understand in any cross-study comparison and 2) independent replication of differential expression results for a disease phenotype would add value to the field. In the case of DCM, by harmonizing data between studies in HeartMap and focusing on genes with consistent effects across all three studies, we can identify a more robust set of genes and pathways that are dysregulated in DCM.

Finally, our analysis of the fibroblast populations demonstrated several sub-populations of interest, namely activated fibroblast groups that were present in specific cardiomyopathies. Activated fibroblasts are a unique population as they are involved in dynamic extracellular matrix remodeling and present an opportunity for therapeutic intervention. Most activated fibroblasts are a result of chronic scar formation with re-purposed functions, such as maintaining structural integrity in damaged tissues, modulating inflammation, angiogenesis and phagocytosis of cells in beneficial and detrimental ways. By identifying marker genes and understanding the pathways involved in these unique populations of activated fibroblasts, we are one step closer to understanding which fibroblast sub-populations are ideal candidates for therapeutic intervention.

## Limitations

Aggregating data sets across multiple studies poses many challenges. We could not control for the variability that exists when isolating nuclei and sequencing tissues across multiple lab environments. Although we separately pre-processed the datasets with equivalent transcriptomes and quality controls methods, we could not account for different equipment, sequencing protocols, and nuclei isolation techniques that vary between laboratories.

Notably, the samples we have included in this atlas are primarily white or non-classified ethnicities. The under representation of certain ancestral backgrounds can pose challenges to applying the findings of this study to a broader population.

## Conclusion

HeartMap provides a comprehensive reference for human heart disease exploration. We have systematically re-processed and integrated multiple datasets together to provide a resource to study heart disease across multiple individuals and cell types. We hope this resource paves the way for a more complete understanding of the human heart in health and disease.

## Online Methods

### Ethics approval and consent

Ethics approval and consent for each study is as follows. For refs^7,10,11,29,30^, including unpublished work from our group, research use of tissues was approved by the institutional review boards at the Gift-of-Life Donor Program, the University of Pennsylvania, Massachusetts General Hospital, and the Broad Institute. Written informed consent for research use of donated tissue was obtained from next of kin in all cases. For ref^6^, heart tissues (donors D1–D7 and D11) were processed at Wellcome Sanger Institute (Hinxton, UK) and obtained from deceased transplant organ donors after Research Ethics Committee approval (ref 15/EE/0152, East of England Cambridge South Research Ethics Committee) and informed consent from the donor families. Heart tissues (donors H2–H7) were processed at Harvard Medical School (Boston, Massachusetts, USA) and obtained from deceased organ donors after Human Research Ethics Board approval Pro00011739 (University of Alberta, Edmonton, Canada). Informed consent from donor families was acquired via the institutional Human Organ Procurement and Exchange Program (HOPE). For ref^12^, research use of tissues was reviewed and approved by the ethics boards of the following institutions: Biobank of the Heart and Diabetes Center NRW, University Hospital of the Ruhr-University Bochum, the HELP Program (Mazankowski Alberta Heart Institute) and HOPE program (University of Alberta Hospital), the Cardiovascular Research Centre Biobank at the Royal Brompton and Harefield hospitals and NHS Blood and Transplant, and the Brigham and Women’s Cardiovascular Tissue Bank. For ref^8^, the local ethics committee of the Ruhr University Bochum in Bad Oeynhausen, the RWTH Aachen University, Utrecht University and WUSTL approved all human tissue protocols (no. 220-640, EK151/09, 12/387 and no. 201104172 respectively). The collection of the human heart tissue was approved by the scientific advisory board of the biobank of the University Medical Center Utrecht, The Netherlands (protocol no. 12/387). All patients provided informed consent, and the study was performed in accordance with the Declaration of Helsinki. For ref^9^, the study complied with all relevant ethical regulations and was approved by the Washington University Institutional Review Board (study no. 201104172). All samples were procured and informed consent obtained by Washington University School of Medicine. For ref^5^, all research tissues were obtained after approval from the Research Ethics Committee and written informed consent were provided by families. The following ethics approvals for donors were obtained: D8 and A61 (REC reference 15/EE/0152, East of England Cambridge South Research Ethics Committee); AH1 (DN_A17), AH2 (DN_A18), AH5 (DN_A19) and AH6 (DN_A20) (REC reference 16/LO/1568, London, London Bridge Research Ethics Committee); AV1 (HOPA03), AV3 (HOPB01), AV10 (HOPC05), AV13 (HOPA05) and AV14 (HOPA06) (REC reference 16/NE/0230, North East, Newcastle & North Tyneside Research Ethics Committee). Samples of failing hearts used for validation were obtained under the Research Ethic Committee approval given to the Royal Brompton & Harefield Hospital Cardiovascular Research Centre Tissue Bank (REC reference 19/SC/0257).

### Aggregating datasets

We queried the PubMed database using terms “snRNA-seq or scRNA-seq or single-cell transcript” and “heart or cardiac or cardiology” as well as web-based searches for single-nucleus RNA sequencing studies. We found twenty adult human heart studies through this search process from 2017-2023. Each study focused on one or a variety of diseases and cardiac anatomies. Of these, nine studies met our inclusion criteria including but not limited to: similar study design (e.g., human-derived samples, 10X Genomics kits), individuals ages greater than 18, nuclei-based analysis, and having greater than 10,000 nuclei (**Table 1**). We limited our study to include only nuclei to avoid transcriptomic differences that exists between cell and nuclei-level data.^17^

One dataset by Simonson et al., establishing the relationship between ARVC (arrhythmogenic cardiomyopathy) and non-failing samples, was included from internal data and has not been made publicly available as of the time of this writing. Additionally, studies from Reichart et al. and Kanemaru et al. included sequencing libraries from control samples that were sequenced as part of previous studies. In these cases, the library was only included once in our atlas. In other cases where sample donors were repeated across studies, but separate sequencing libraries were constructed, we kept both sample files but carefully conducted downstream analyses to account for possible over-abundance of samples from one individual.

### Single nuclei sequencing and pre-processing of datasets

Sequencing protocols were relatively consistent across studies and can be found in their respective publications. However, a few differences were notable regarding the single-nuclei purification and 10x genomics library construction. Multiple studies included nuclei purified by fluorescent activated cell/nuclei sorting (FACS/FANS) including Litvinukova et al., Kuppe et al., Koenig et al., and Kanemaru et al. Many of the included datasets used 10x Genomics 3’ version 3 chemistry (3’v3), however Tucker et al. used 3’v2, Litvinukova et al. used a combination of 3’v2 and v3, and Koenig et al. used 5’v1.1.

We ensured equivalent pre-processing of the raw FASTQ files across studies by using a similar transcriptomic reference and software packages (**Supplemental Table 7**). FASTQs were trimmed for the template switch oligo (TSO) and homopolymers (A_30_, C_30_, T_30_, G_30_) with *cutadapt* prior to alignment when possible. We used pre-mRNA GRCh38 2020-A as our reference transcriptome and 10X Genomics CellRanger v4.0.0 for transcriptome alignment and mapping when possible. In the case of Koenig et al., which used 5’v1.1 with an extended R2 length, we trimmed R2 down to 91BP to be consistent with 10X Single Cell 3’ v3 read lengths used in most other studies. We confirmed that this reprocessing resulted in highly correlated gene expression to published counts **(Supplemental Figure 21)**. scR-Invex (https://github.com/getzlab/scrinvex) was used for assigning intronic, exonic and junction spanning reads.^31^ CellBender v2.0.0 (https://github.com/broadinstitute/CellBender) and Scrublet were used for additional empty droplet detection and doublet detection, respectively.^32,33^

Nuclei quality control (QC) was conducted by sub-setting nuclei to those kept in the original published datasets to maintain consistency with publications. In some rare cases, CellBender removed all gene counts in a nucleus and these nuclei were excluded. Additionally, we performed new QC on the data from Tucker et al. using the process described in the published Chaffin et al. and Simonson et al. studies.

In one study (Kuppe et al.), similar pre-processing of datasets was not possible because raw FASTQ files were not provided by the study authors at the time of data collection. In this case, the provided count data used an unknown GRCh38 transcriptome reference with CellRanger v6.0.2 used for alignment. We confirmed that the genes represented in this provided dataset were identical to those in our internal datasets included in our atlas. CellBender and Scrublet was run on the full count data to generate a dataset as equivalent to the other studies as possible. Additionally, we excluded samples from this study (CK161, CK369, CK376) due to poor quality and low UMI counts.

### Metadata Harmonization

Metadata was provided for each study in the supplemental information of each publication. However, meta data varied across studies. Few studies consistently reported data such as age, biological sex, race, ethnicity, weight/body mass index, heart weight, and left ventricular ejection fraction.

X and Y chromosome gene counts were calculated for each sample to determine the transcriptional sex for each sample, using *DDX3Y* for the Y chromosome and *XIST* for the X chromosome. There were a few discrepancies in samples between the metadata sex assignment and chromosome-count based sex analysis. These samples are from the Litvinukova et al. (H0015_RV), Reichart et al. (BO_H40_LVW), and Kanemaru et al. (HCAHeartST13146207, HCAHeartST13146208). In the case of H0015_RV and BO_H40_LVW, the respective studies had multiple samples from the same donor. We confirmed that these two samples did not share the same genetic fingerprint as the other samples from their respective donors using CrosscheckFingerprints^34^ from picard (https://github.com/broadinstitute/picard). Samples with discordant phenotypic sex and X/Y chromosome counts from the snRNA-seq data were excluded from the atlas. All analyses requiring sex information used the chromosome-derived sex assignment, unless otherwise noted.

Additionally, we excluded samples from multiple studies with ages less than or equal to 19 to ensure only adult samples were include in our atlas. This age range was selected because Reichart et al. used age ranges such as 0-19 to describe non-adult individuals. The studies with these excluded samples are from Reichart et al. (H05, H59, H73, H79, H80, H81, DO1, H84, IC_H02_LV0) and Koenig et al. (TWCM-11-93, TWCM-13-181, TWCM-13-198, TWCM-13-235). Furthermore, when calculating age for our summary statistics, we treated ages presented as ranges as the median value (e.g. age in years 60-65 became 62). Koenig et al. had one individual with missing age in published metadata (TWCM-11-3). Age was not included as a covariate in our differential expression testing due to the labelling of ages using ranges as opposed to discrete values.

### Batch-correction and benchmarking

We selected several batch-correction methods from the top-performing methods identified in a previous study of benchmarking cell and nuclei integration on a large atlas.^17^ These methods included Harmony, Scanorama, Single-cell variational inference (scVI), and Single-cell annotation using variational inference (scANVI).^13–16^ Specifically, scANVI required prior cell type assignments to run thus we generated crude labels for scANVI using the Leiden clustering algorithm on a non-batch corrected neighborhood graph with resolution of 1.0 and assigning cell types based on known canonical cell type genes. Each method had various tunable parameters such as whether to estimate highly variable genes (HVG) by batch or across all nuclei, and the number of principal components (PC) or size of the latent space (LS). For each method we tested multiple combinations of these parameters such as inclusion of the top 2000 HVGs across all nuclei or by batch (i.e., donor of origin) using methodology from Seurat V3^35^, PCs of either 50 or 100, and LS of either 10 or 25. This resulted in four versions for each batch-correction method for a total of sixteen variations of batch-correction. Each unique method was applied to a subset of the atlas (619,918 nuclei or one fourth of the HeartMap atlas) that was representative of the proportion of studies from the overall atlas. The batch key for integration was set as *individual* (representing the donor of origin) with an additional co-variate key set to *study* to reduce technical noise across samples and datasets. *Individual* was selected as our co-variate for alignment as this appears to be a primary cause of batch-effects in cardiac cells, as seen in Tucker et al. (Supplemental Figure 2).^7^

Batch correction evaluation and comparison between the sixteen parameterizations of correction methods was performed using existing integration benchmarking metrics.^17^ The benchmarking metrics included important measures of biological conservation and batch correction at the principal component analysis (PCA) embedded space and k-nearest neighbors graph space levels. Methods that measured biological conservation included normalized mutual information (NMI), adjusted Rand index (Ari), isolated and non-isolated average silhouette width (ASW), cell cycle conservation, cell-type local Simpson’s inverse index (cLISI), and F1 score. Methods that measured batch correction included graph connectivity, integration local Simpson’s inverse index (iLISI), batch ASW, and principal component regression. Metrics were calculated using *individual* and *study* as the covariate keys to measure performance across these two variables (**Supplemental Figure 1**). After the metrics were calculated for each of the sixteen batch-correction method, we calculated a scaled score for ranking each metric performance across the sixteen methods calculated as reported in Luecken et al. 2022.^17^

The final score and each partial score were the objective measures used to determine the batch-correction metric applied to the entire atlas. In cases when scores for batch-correction and biological conservation were extremely close, we visually assessed the mixing of nuclei in UMAP plots colored by individual, study, and cell type labels to determine the optimal batch-correction method.

### Map Aggregation, cell type annotation, and marker genes

Single-nuclei RNA sequencing map aggregation was performed using scanpy v1.9.3 unless otherwise stated.^36^ The atlas map was constructed using the post-QC nuclei that remained after sub-setting to published data as previously described. Additionally, we assigned cell type labels based on the expression of canonical markers of known cell types using Leiden clustering, as described previously. Based on the results of the top-performing batch-correction method, scANVI was used to harmonize the atlas after calculating highly variable genes across all nuclei and using a latent space of 25 latent variables. We used the rough cell type labels as input into scANVI for batch correction. A neighborhood graph was constructed using these 25 latent variables with sc.pp.neighbors() using 15 neighbors and *cosine* distance. UMAP was constructed with the effective minimum distance between embedded points set to *min_dist = 0.2*. Leiden clustering was employed at 2.0 resolution to help determine cell type composition of the post-batch corrected atlas.

Marker genes were calculated for each global cluster using an area under the receiver operating characteristic curve (AUC). The AUC was computed for each gene comparing the log-normalized expression at the nuclei-level in each cluster against all other clusters. A final list of marker genes for each cluster was ascertained using an AUC threshold > 0.6. Marker genes for each cluster verified the existence of canonical genes that defined each global cluster by cell type.

### Sample-level principal component and UMAP analysis

PCA was performed on the sample-level to identify high level transcriptional differences based on phenotypic data. In this case, sample was defined by a unique identifier consisting of the individual name + study of origin + anatomy. For example, a sample from the Chaffin et al. dataset would be listed as P1304-Chaffin-LV. We summed the CellBender-derived gene counts across all nuclei per sample, treating the data as a bulk experiment. Mitochondrial genes, ribosomal genes and genes expressed in less than or equal to 1% of nuclei were removed. Remaining genes were normalized with DESeq2 with a design ∼ disease + sex and genes with less than 10 total counts were removed.

A variance stabilizing transformation was applied with *vst()*. Principal components were calculated with *prcomp()* in *R*. Technical effects were removed by adjusting for the *study* covariate. PCA was run on the top 500 highly variable genes using 30 PCs, with the exception of epicardial cells which used only 26 PCs due to the number of samples available. A neighborhood graph was constructed with *sc.pp.neighbors()* using 15 neighbors and *cosine* distance. A UMAP representation was constructed using *min_dist* = 0.2. When calculating PCA and UMAP by cell type, our model design was the same as above, hence we removed atrial cardiomyocytes which were from only control samples.

### Pairwise differential expression analysis based on disease status and sex-assignment by cell type

Differential expression between all pairwise disease comparisons was performed on a cell type basis. The analysis was limited to only the ventricular anatomies (LV, RV, AX and IVS) as non-ventricular samples only came from control samples. We removed studies including: Litvinukova et al., Tucker et al., and Kanemaru et al., because these only had control hearts which may not contribute meaningfully to comparisons between diseases. Additionally, the disease CAD-HF was excluded as it was only included in one sample in the Kuppe et al. study. Testing was performed using the limma-voom framework. Nuclei were summed for a sample, defined by a unique identifier consisting of the individual name + study of origin + anatomy, if there were > 25 nuclei for the given cell type. Genes in < 0.5% of nuclei in all disease groups were removed. Next, a differential expression model was performed using the limma-voom framework using DESeq2 normalization.^37,38^ We used the model *Expr* ∼ 0 + disease + study + anatomy + sex, with a *duplicateCorrelation()* effect fit for the individual donor of origin to control for the fact that some individuals are represented across multiple studies or anatomies. We then extracted 15 pairwise disease contrasts for each cell type if there were at least 3 samples in the two groups being compared. Differential expression was performed on CellBender counts as primary analysis, as well as CellRanger counts as a sensitivity analysis. A background heuristic, as described in Simonson et al. 2023, was calculated using all nuclei from each disease group being compared. Only genes with a background heuristic based on nuclei from group 1 and group 2 < 0.5 were included. Additionally, only genes with Benjamini-Hochberg corrected p-values < 0.05 with both CellBender and CellRanger that were expressed in at least 0.5% of nuclei from one comparator group in a given contrast were deemed significant.

Differential expression between sex-assignment was performed on a cell type basis. The analysis included all anatomical regions and studies; however, we again excluded the disease CAD-HF. Testing was performed as above, and we extracted 1 pairwise sex-assignment contrast for each cell type if there were at least 3 samples each in the female and male groups. Only genes with a background heuristic based on nuclei from group 1 and group 2 < 0.5 were included. Additionally, only genes with Benjamini-Hochberg corrected p-values < 0.05 with both CellBender and CellRanger were deemed significant.

### Comparing DCM and NF across studies

To understand the consistency of differential expression tests across studies, we focused on the comparison of DCM to non-failing control hearts as DCM was captured in 3 distinct studies (Chaffin et al., Reichart et al., and Koenig et al.). Within each study, we ran individual DE tests between DCM and non-failing controls by cell type using the reprocessed data in our atlas. In the Chaffin et al. re-processed data, we included 50 samples (27 individuals total, 11 DCM and 16 controls) from the LV as this was the only anatomy included in the publication. We summed across repeated samples per individual so that each individual was only represented once in the DE test. In the Reichart et al. re-processed data, we included 111 samples (50 individuals total, 44 DCM and 6 controls) and all anatomies studied. Of note, the original publication used 12 additional controls derived from Litvinukova et al. which we excluded here as they were not sequenced as part of the Reichart et al. study. To create pseudo-bulk expression, we summed across the combination of individual + anatomy. In the Koenig et al. re-processed data, we included 34 samples (34 individuals total, 11 DCM and 23 controls) from AX samples as this was the only anatomy included in the publication. As there were no duplicate samples per individual, we summed across individual. For Chaffin et al. and Koenig et al., differential expression testing was performed as previously described using the limma-voom framework with the model ∼ 1 + disease + sex, requiring a minimum of 25 nuclei and 3 samples per cell type group. For Reichart et al., because multiple anatomies were included, we used the model ∼ 1 + disease + sex + anatomy with a duplicateCorrelation() effect for individual to control for the fact that some individuals are represented across multiple anatomies. Only genes expressed in at least 0.5% of nuclei in either DCM or control for a given study were included.

When possible, we compared published and re-processed estimates for each study (**Supplemental Figure 5A,B**). As Reichart et al. did not report a direct DCM to control comparison, we could not check for consistency with our effect estimates. To assess heterogeneity in our re-calculated effect estimates between DCM and control across studies, we estimated the I^2^ metric using the *metafor* package in R (**Supplemental Figure 5C**).

### Sub-clustering analysis

Global clusters were collapsed into cell types based on canonical gene markers. If there were two or more global clusters with canonical genes mapping to a single cell type, we aggregated them together for downstream analysis. To create each cell type-specific map, we used scVI, instead of scANVI, for batch-correction with HVG calculated across all nuclei and a latent space of 25 because there was only one cell type for the basis of the integration.

Once a cell type-specific map was created, we applied a different Leiden clustering strategy than previously described to overcome sub-cluster gene-expression similarity that was less apparent on the global atlas. To select the optimal clustering resolutions for each cell type-specific map, we generated Leiden clustering resolutions from 0.1 to 3.0, and at each resolution we pulled out genes from each sub-cluster with AUC > 0.55 and LogFC > 1.5 compared to all other sub-clusters combined. If there were more than 50,000 cells in the cell type, we stopped at the first resolution where greater than or equal to 5 sub-clusters had less than or equal to 10 genes. If there were less than 50,000 cells in the cell type, we stopped at the first resolution where at least 1 sub-cluster had less than or equal to 10 genes.

Marker genes for final sub-clusters were computed using the AUC-derived test as above as well as a limma-voom-derived test based on sample level aggregation. In addition to the contrasts that differentiated each sub-cluster versus all other sub-clusters in the limma-voom differential expression analysis, we also computed all pairwise contrasts for each sub-cluster versus every other sub-cluster to further define sub-cluster differences and choose marker genes for validation. The marker genes for validation were selected based on AUC > 0.6 and high logFC estimates > 1.5 relative to other sub-clusters.

### Compositional analysis

To identify compositional differences across and within cell types, we applied the Bayesian-based model from scCODA v0.1.9.^39^ For this analysis, a reference cell type is required that has a nearly constant abundance across samples. For our analysis we manually set the reference cell type to the global cluster or sub-cluster with the least dispersion across samples. For each test we ensured the acceptance rate was > 35%. Our model included the covariates: disease, anatomy, study, and sex.

### Gene set enrichment analysis

Pathway analysis was performed on the sub-clusters within each cell type using the open source Bioconducter R packages fgsea^40^ and clusterProfiler^41^ (release 3.18) and the biological processes gene ontology (GO) terms.^42^ For the fgsea analysis, genes for each sub-cluster were ranked based on the t-statistic from the limma-voom based differential expression test. Only GO terms with greater than 15 and less than 500 genes were considered. Ontologies with a Benjamini-Hochberg adjusted p-value <0.05 and normalized enrichment score >2 were considered significant. For the clusterProfiler analysis, enrichment tests were calculated for GO terms and KEGG pathways based on hypergeometric distribution. Genes from each sub-cluster were selected based on a Benjamini-Hochberg adjusted p-value < 0.05 and logFC > 0 from the limma-voom based differential expression test. Ontologies with a corrected p-value < 0.1 and logFC > 0 were significant.

### Photobleaching, RNA scope, and Immunofluorescent Validation

Fresh frozen tissue sections of 10 um were fixed in 4% PFA for 30 min at RT and washed three times in PBS. Slides were then placed in trays in ice cold PBS and photobleached for 48 h (protocol modified from Sun et al.).^43^ Slides were placed in a large ice bucket containing ice in a black 395 mm high bin within a cold room. Slides placed in PBS to prevent dehydration were placed on top of ice packs inside the bin. A 300-watt full spectrum grow light (Platinum LED Lights, catalog number P300) was placed on top of the bin and turned on with a cloak to cover the bin to prevent UV radiation. Ice, ice packs, and cold PBS were replaced at the beginning and end of each day to keep the slides cold. After 48 h, RNAscope was carried out on the 10 μm fresh frozen tissue sections using the RNAscope Multiplex Fluorescent Reagent Kit v2 using manufacturers protocols. As there were two populations of activated fibroblasts that we were interested in, we used POSTN which is a canonical gene specific to activated fibroblasts. We also identified genes that were highly selective to our activated fibroblast subclusters compared to other fibroblast cell populations (AUC >0.6, log fold-change >1). As these sub-populations are rare, we aimed at validating on cardiac tissue with estimated > 3 percent of nuclei expression in fibroblasts. We selected cardiac tissue from three LV samples for each of the disease groups (inherited disease, ischemic disease, and non-failing controls) used in our prior studies (see Supplemental Table 6). However, we limited our TNC+POSTN+ validation to samples with ischemic cardiomyopathy as no acute myocardial samples were available. Non-failing control samples were selected based on a low, medium, and high TNC+POSTN+ fibroblasts expression profile relative to total fibroblasts using computational prediction of tissue expression levels. The percentage of activated fibroblast populations in the computational prediction and experimental validation were calculated relative to the total number of POSTN+ cells. For example, the percentage of TNC + POSTN+ cells was calculated by totaling the number of cells expressing both TNC and POSTN and dividing the sum by the number of cells expressing POSTN. The probes used for this study were hs-POSTN (409181), hs-COL22A1 (818811-C2) and hs-TNC (420771-C3). Slides were mounted with Prolong Gold and imaged on a Leica SP8. For quantification, tissue sections were tiled at 20x magnification. For representative images 1 um z stacks were taken through the tissues on 20x.

### Imaging Analysis

Each individual channel from individual tiled images were exported and data was analyzed using the cell counter tool in Image J (version 1.54f). Individual channels for each image were stacked and using the cell counter/point tool positively stained cells were counted on each channel. Using the nuclei channel as a guide, double or triple positively stained cells were identified.

## Acknowledgements

We would like to thank the individuals who provided the samples used in this study. YD conducted work supported by the Sarnoff Cardiovascular Research Foundation. P.T.E. is supported by grants from the National Institutes of Health (RO1HL092577, RO1HL157635, R01HL177209), from the American Heart Association (961045), from the European Union (MAESTRIA 965286) and from the Fondation Leducq (24CVD01).

## Supplemental Figures

**Supplemental Figure 1.**
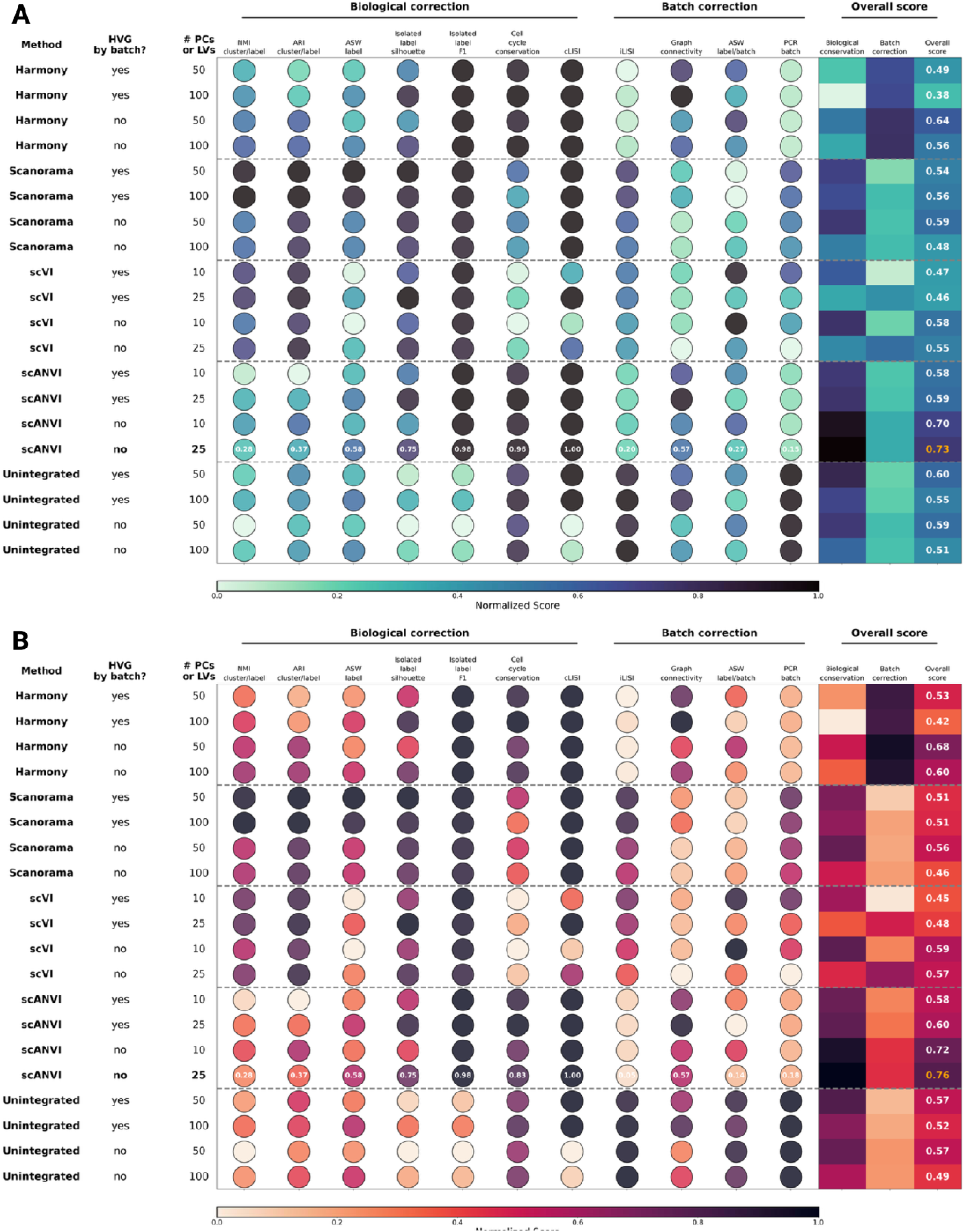
Batch correction benchmarking metrics used to assess the overall scores of the integration methods based on the “individual” and “study” covariate keys. **A, B,** Scoring was conducted using “individual” (**A**) and “study” (**B**) as the covariate keys. Columns keys are as follows: Method = Batch-correction method used, HVG by batch? = using the top 2,000 highly variable genes across each batch, instead of across all nuclei, for batch-correction, # PCs = number of principal components, # LVs = number of latent variables. A normalized scale bar is provided to show how the scores compare to each other (see Methods).

**Supplemental Figure 2.**
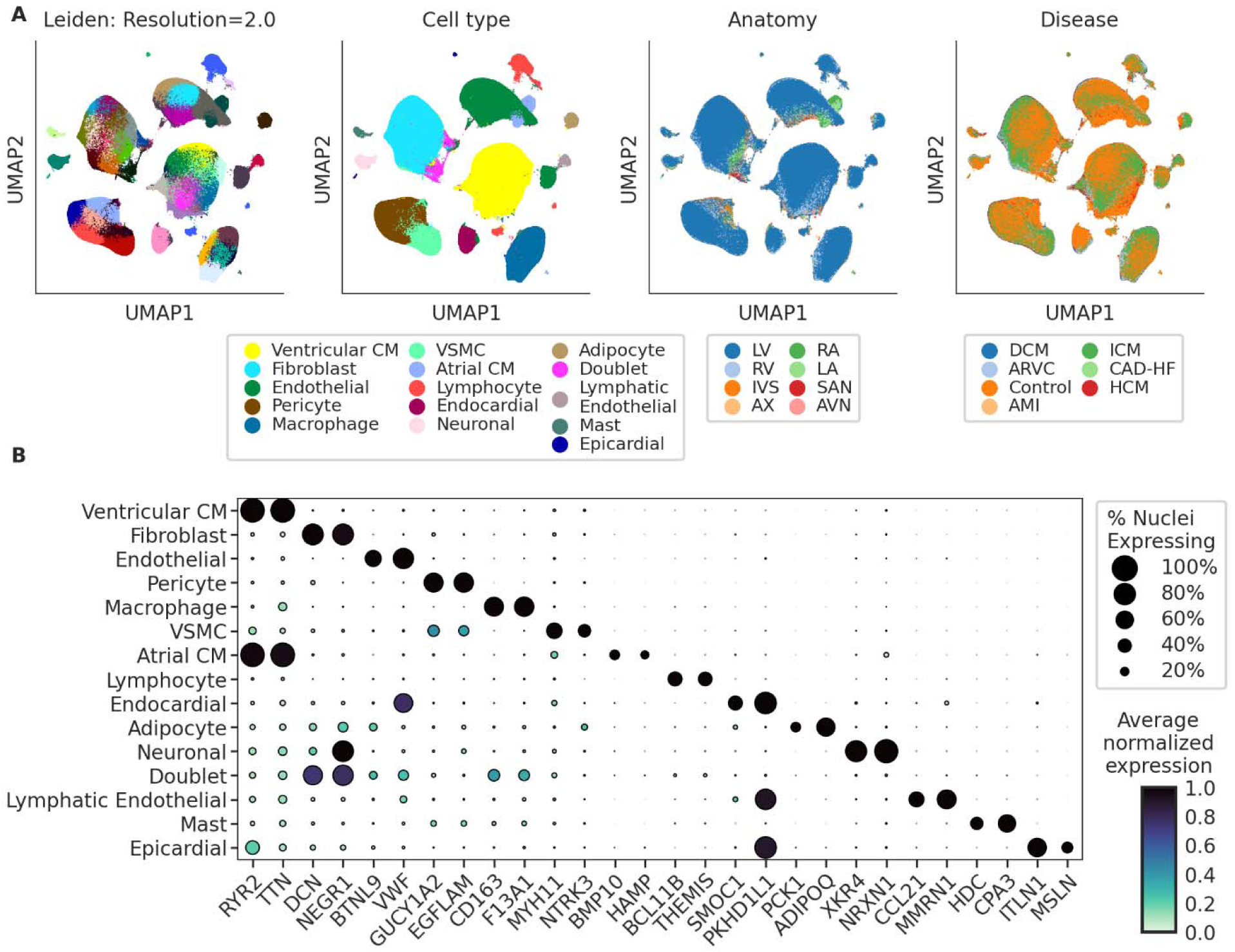
Global aggregation of 2.4 million nuclei integrated across individual and study using scANVI. **A**, UMAP representations of the atlas revealing common and rare cell types. Nuclei are colored by Leiden cluster at resolution 2.0 (left most), cell type (collapsing Leiden clusters), anatomical regions, and disease phenotype (right most). **B**, Dot plot depicting the expression of the top marker genes for each global cell type by highest nuclei-level estimate logFC that are protein-coding and have an AUC-score > 0.6. The size of each point represents the percent of nuclei in the given cluster expressing the gene at non-zero levels and the shading represents the average log-normalized expression of each gene in each cluster, normalized across clusters using min-max normalization.

**Supplemental Figure 3.**
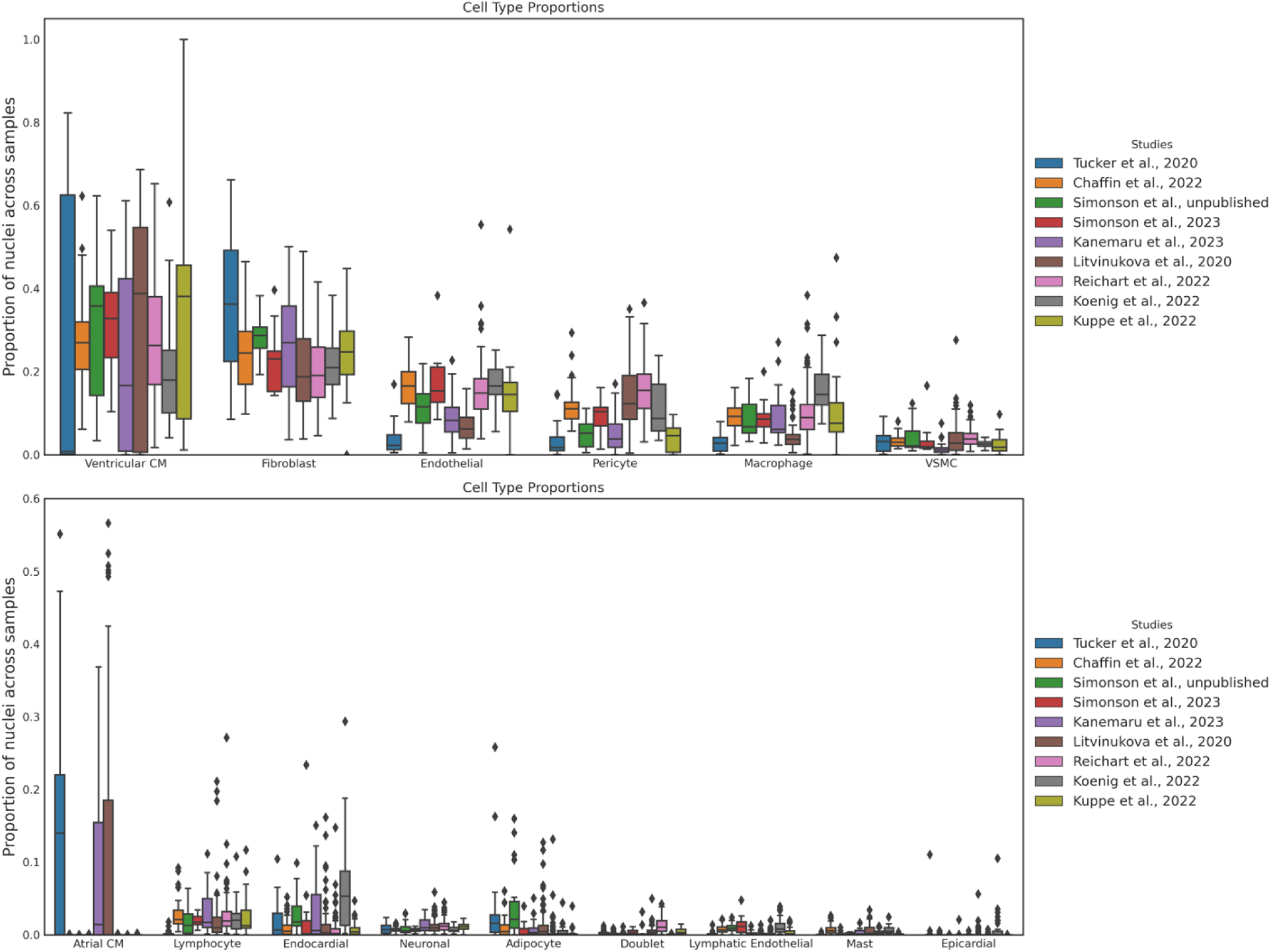
Cell type composition by study. Box and whisker plots show the distribution of cell type proportions, with the center of the box being the median and lower and upper bounds being the 25^th^ and 75^th^ percentiles, respectively. Outliers are shown as points in the box plots. Sample is considered as a unique combination of “study” + “individual” + “anatomy” (see Methods).

**Supplemental Figure 4.**
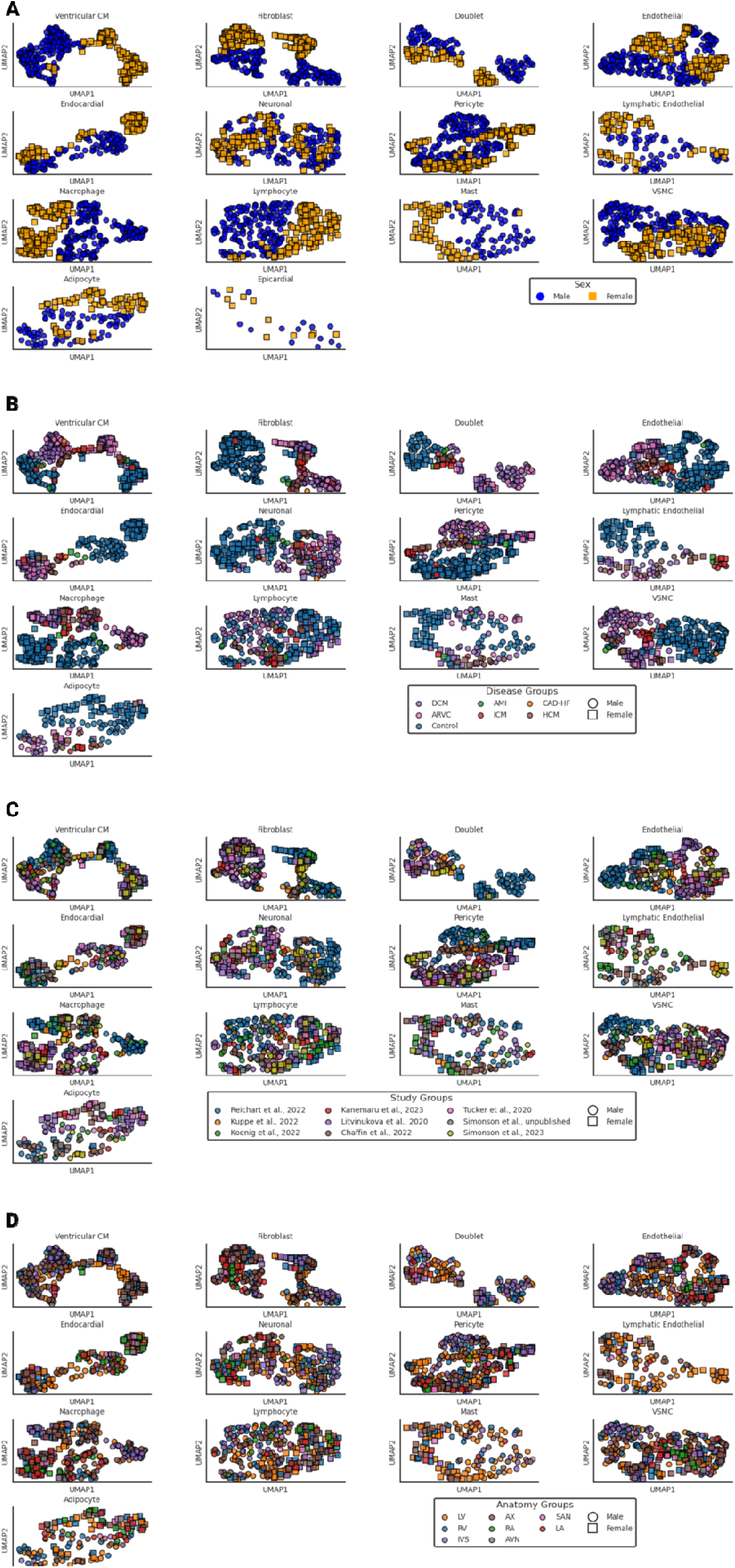
UMAP by cell type based on covariates. **A, B, C, D,** Plots shows the UMAP representations colored by sex-assignment (**A**), disease (**B**), study groups (**C**), and anatomical groups (**D**). Each point on the plot represents one sample. CM, cardiomyocyte; VSMC, vascular smooth muscle cell.

**Supplemental Figure 5.**
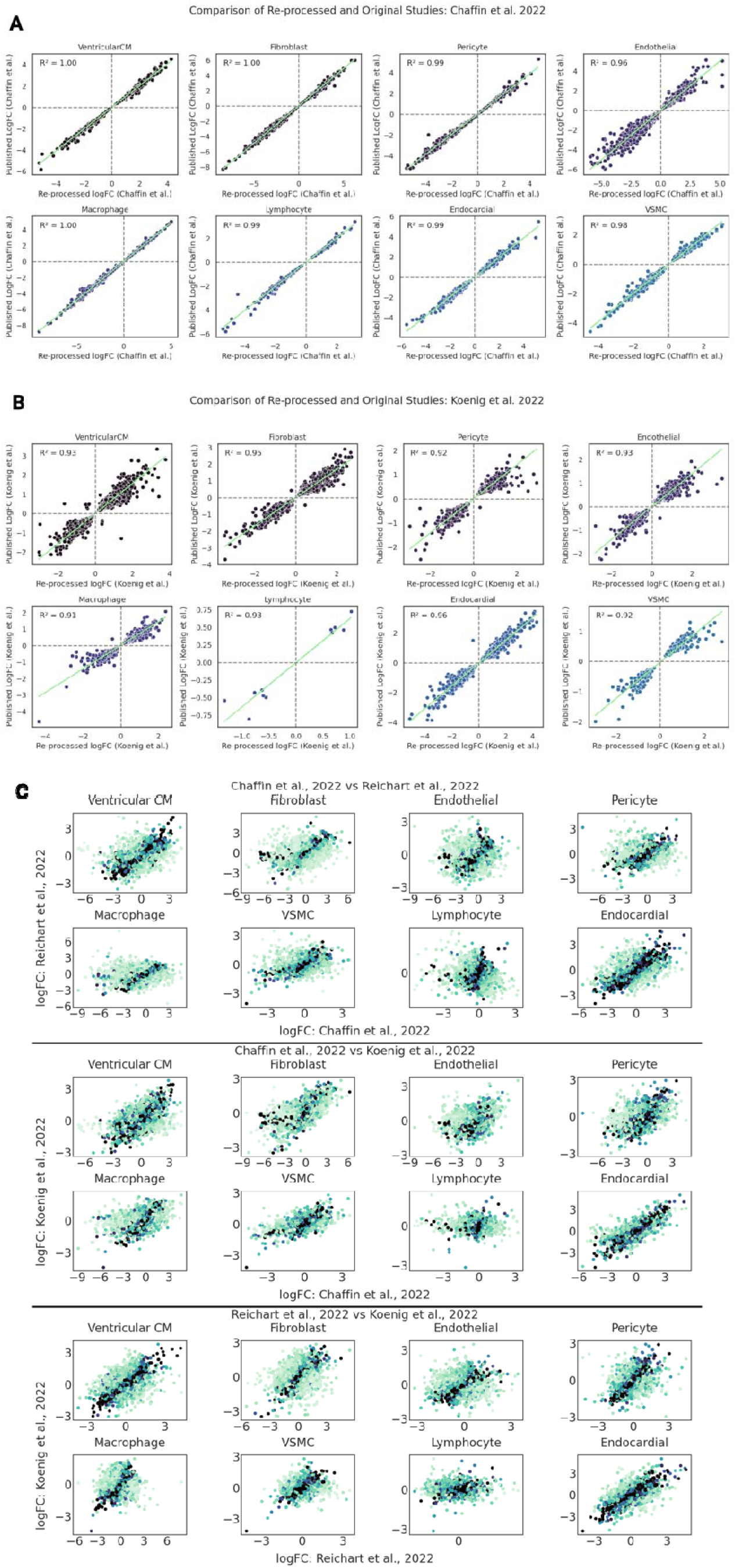
Correlation plots for effect estimates (LogFC) between studies and DCM vs NF control analysis. **A, B,** Effect estimate (LogFC) correlations are shown between genes from published studies and the reprocessed studies in HeartMap. Plots for each study showing LogFC correlation based on the computed (Re-processed) and published values are presented for Chaffin et al., 2022 (**A**), and Koenig et al., 2022 (**B**). Reichart et al., 2022 did not have published effect estimates and was not included in this figure (see Methods). Only genes with a significant differential expression result in the re-processed analysis are included. **C,** Correlation plots are shown comparing each re-processed study’s effect estimates across cell types in the DCM vs NF control analysis. Each point in the plot (**C**) represents a shared gene between both studies. Genes are colored based on the concordance of effect estimates using the I2-coefficient, a measure of heterogeneity between studies. CM, cardiomyocyte; VSMC, vascular smooth muscle cell.

**Supplemental Figure 6.**
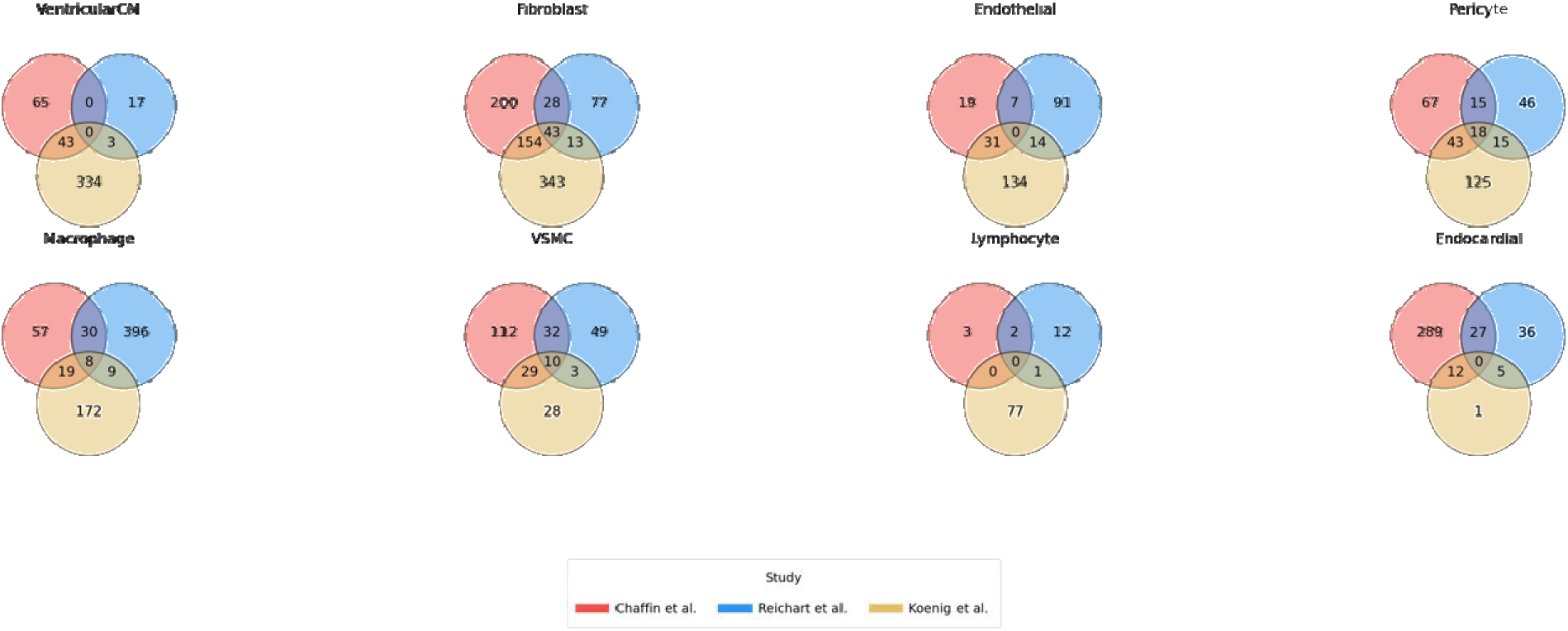
Venn diagrams demonstrating the overlap of pathways in DCM vs NF control samples across the multiple re-processed studies. Pathways for each study, Chaffin et al., 2022, and Koenig et al., 2022. Reichart et al., 2022 were derived using gene-set enrichment analysis, with adj. p-val < 0.05. CM, cardiomyocyte; VSMC, vascular smooth muscle cell.

**Supplemental Figure 7.**
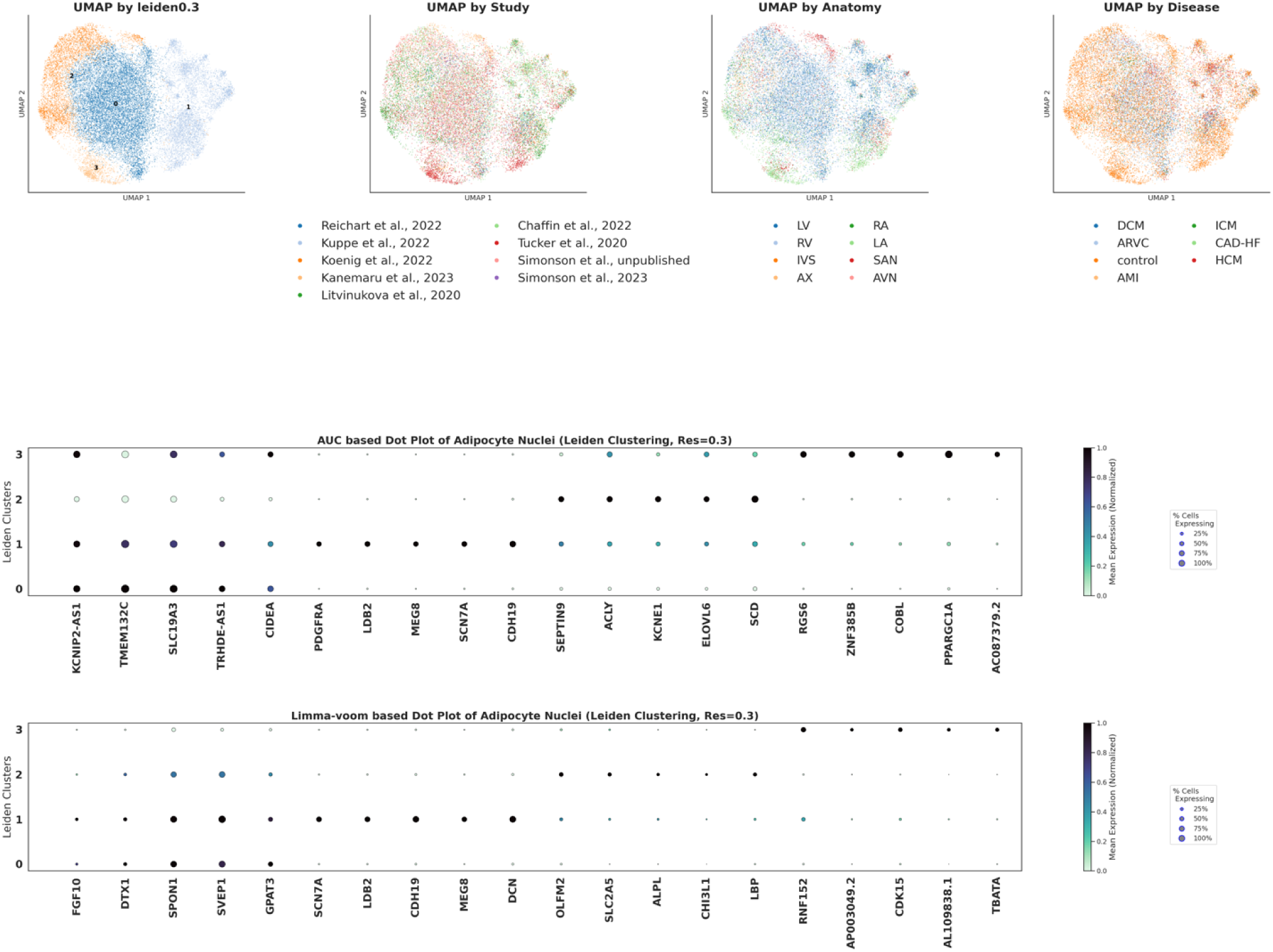
UMAP and Dot plot for HeartMap Adipocyte nuclei. UMAPs are depicting Leiden-clustering resolutions, study, anatomy, and disease, where each point represents a nucleus, and colors differentiate the co-variate. Dot plots show the top AUC-score based and limma-voom based markers by log fold change for each Leiden cluster (see Methods).

**Supplemental Figure 8.**
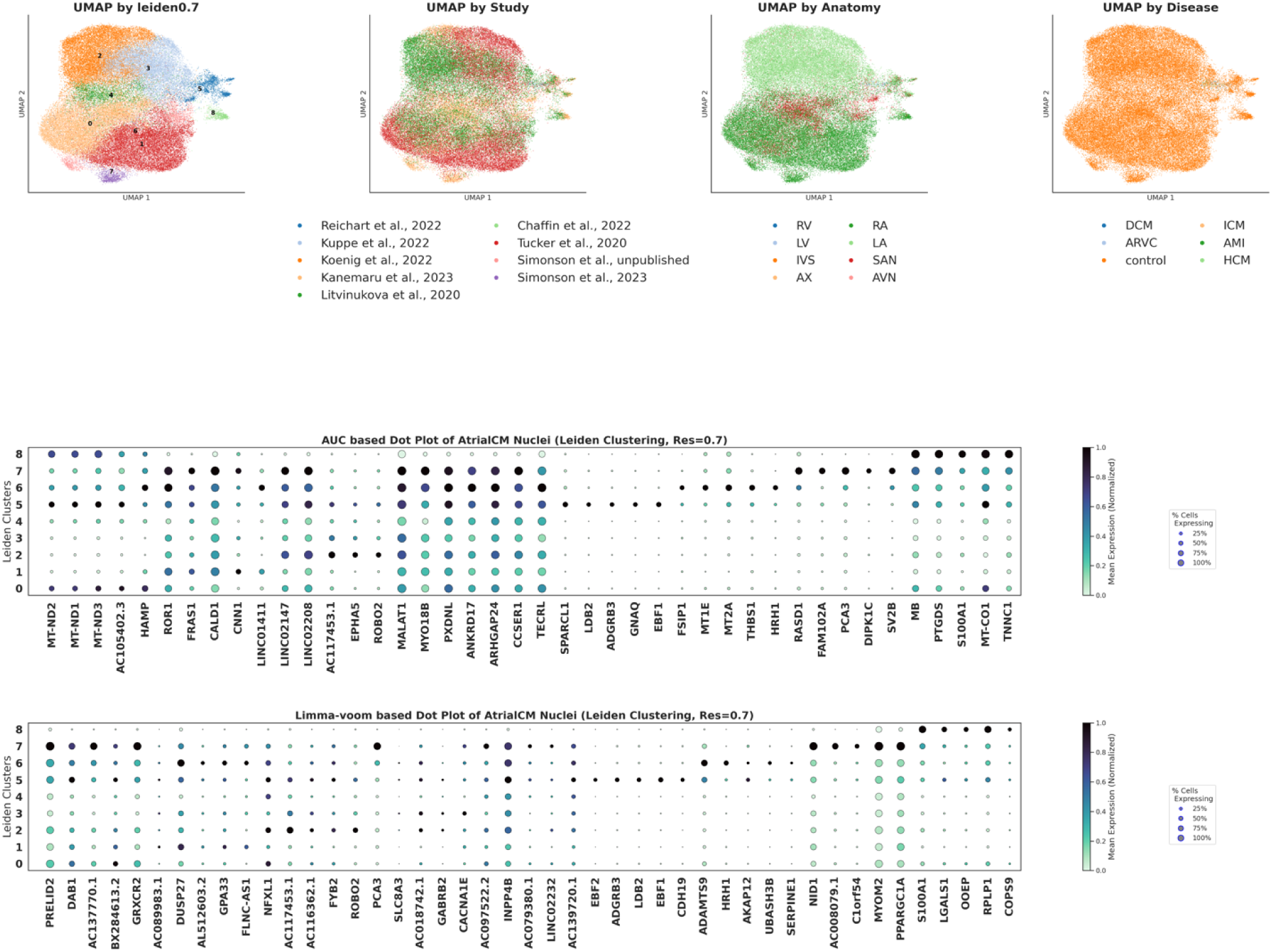
UMAP and Dot plot for HeartMap Atrial Cardiomyocyte nuclei. UMAPs are depicting Leiden-clustering resolutions, study, anatomy, and disease, where each point represents a nucleus, and colors differentiate the co-variate. Dot plots show the top AUC-score based and limma-voom based markers by log fold change for each Leiden cluster (see Methods).

**Supplemental Figure 9.**
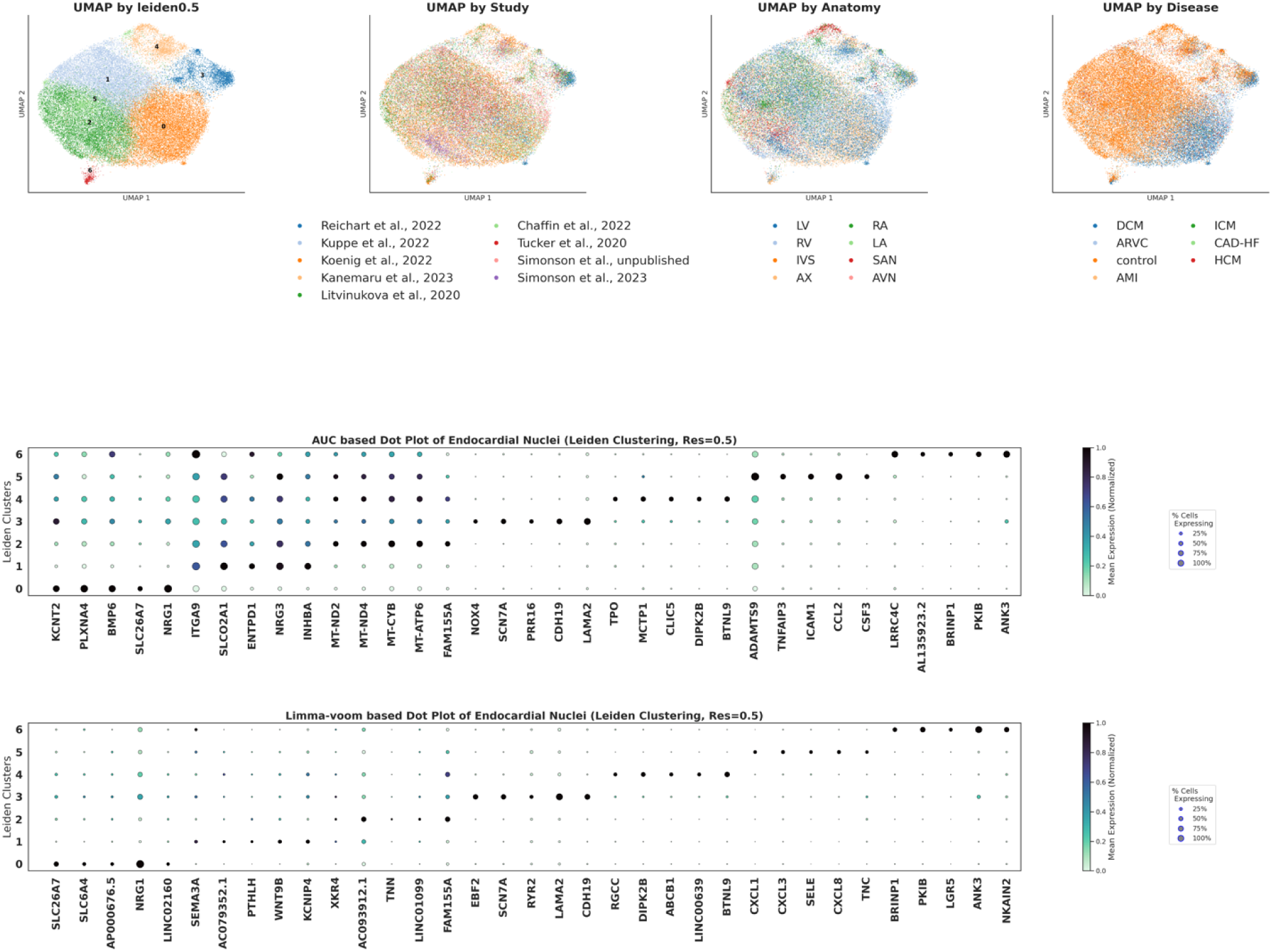
UMAP and Dot plot for HeartMap Endocardial nuclei. UMAPs are depicting Leiden-clustering resolutions, study, anatomy, and disease, where each point represents a nucleus, and colors differentiate the co-variate. Dot plots show the top AUC-score based and limma-voom based markers by log fold change for each Leiden cluster (see Methods).

**Supplemental Figure 10.**
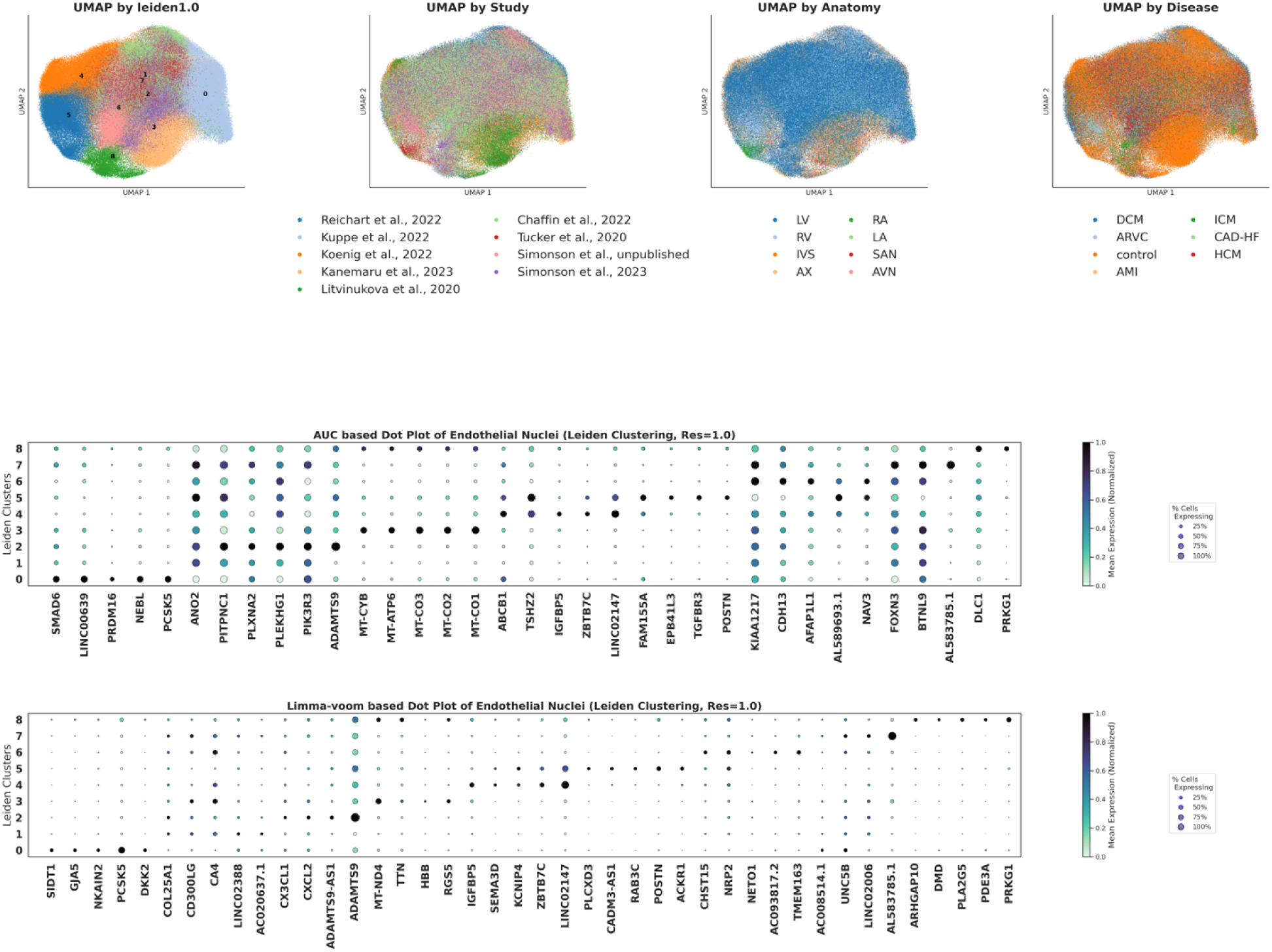
UMAP and Dot plot for HeartMap Endothelial nuclei. UMAPs are depicting Leiden-clustering resolutions, study, anatomy, and disease, where each point represents a nucleus, and colors differentiate the co-variate. Dot plots show the top AUC-score based and limma-voom based markers by log fold change for each Leiden cluster (see Methods).

**Supplemental Figure 11.**
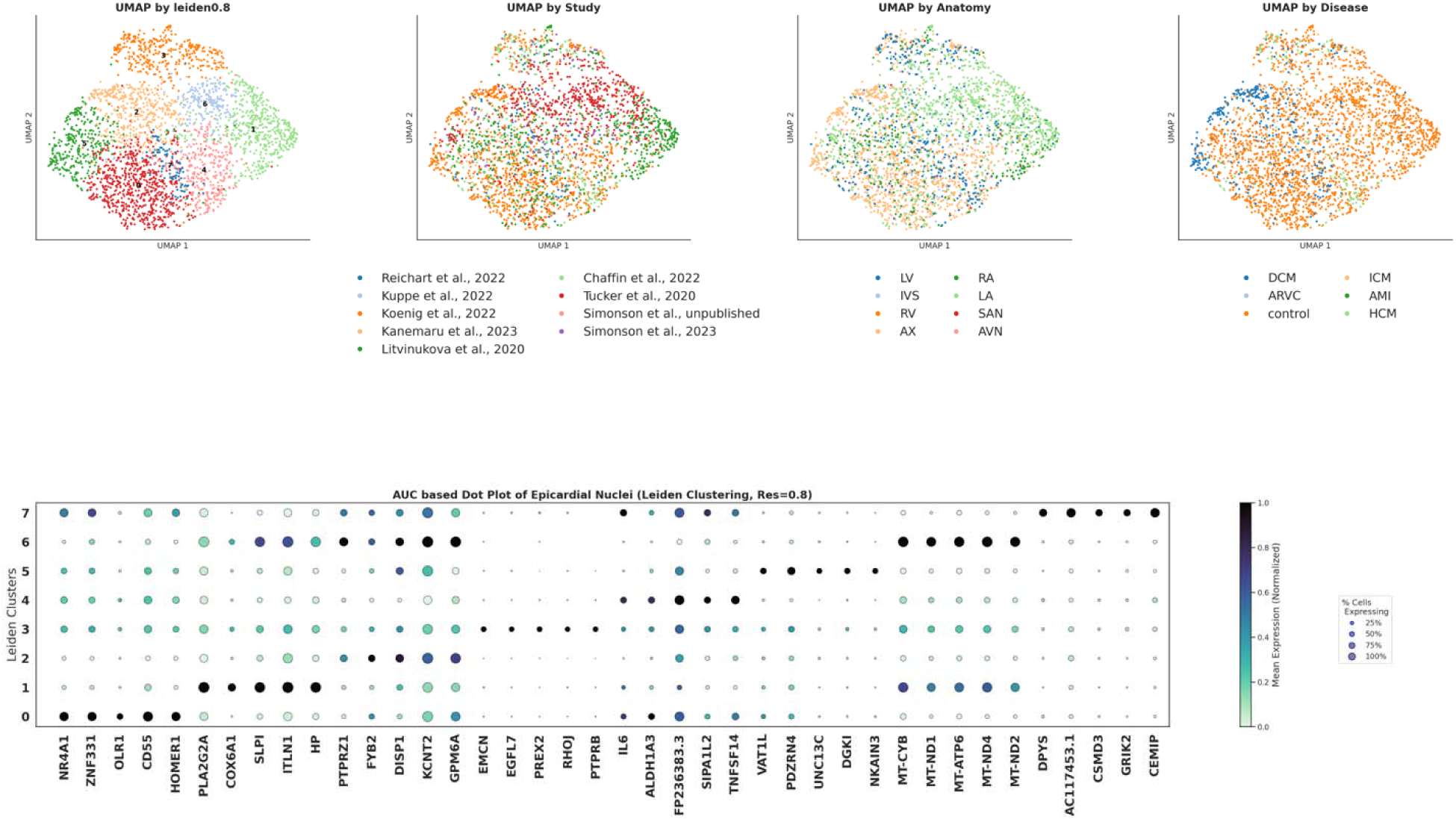
UMAP and Dot plot for HeartMap Epicardial nuclei. UMAPs are depicting Leiden-clustering resolutions, study, anatomy, and disease, where each point represents a nucleus, and colors differentiate the co-variate. Dot plots show the top AUC-score based and limma-voom based markers by log fold change for each Leiden cluster (see Methods).

**Supplemental Figure 12.**
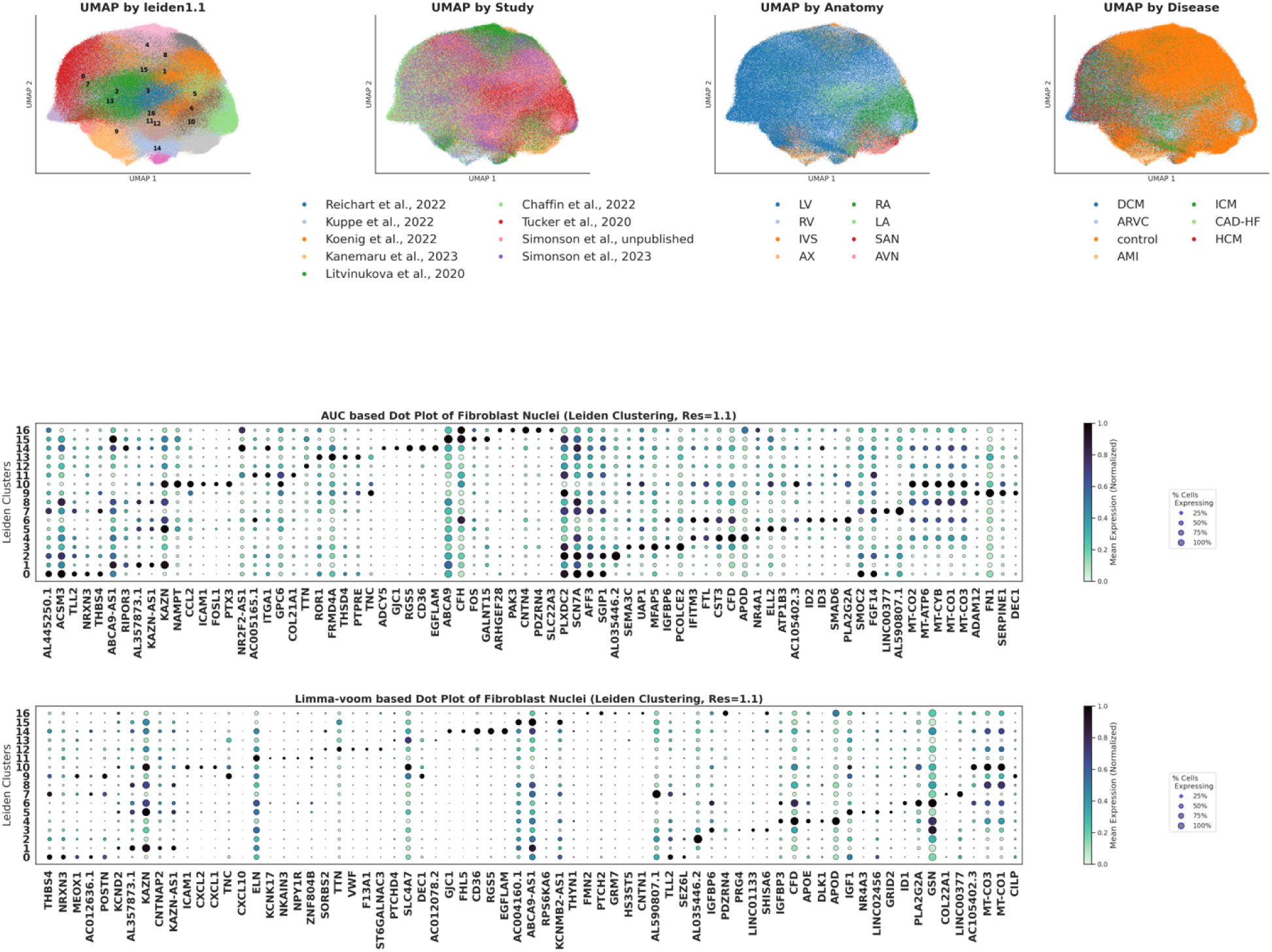
UMAP and Dot plot for HeartMap Fibroblast nuclei. UMAPs are depicting Leiden-clustering resolutions, study, anatomy, and disease, where each point represents a nucleus, and colors differentiate the co-variate. Dot plots show the top AUC-score based and limma-voom based markers by log fold change for each Leiden cluster (see Methods).

**Supplemental Figure 13.**
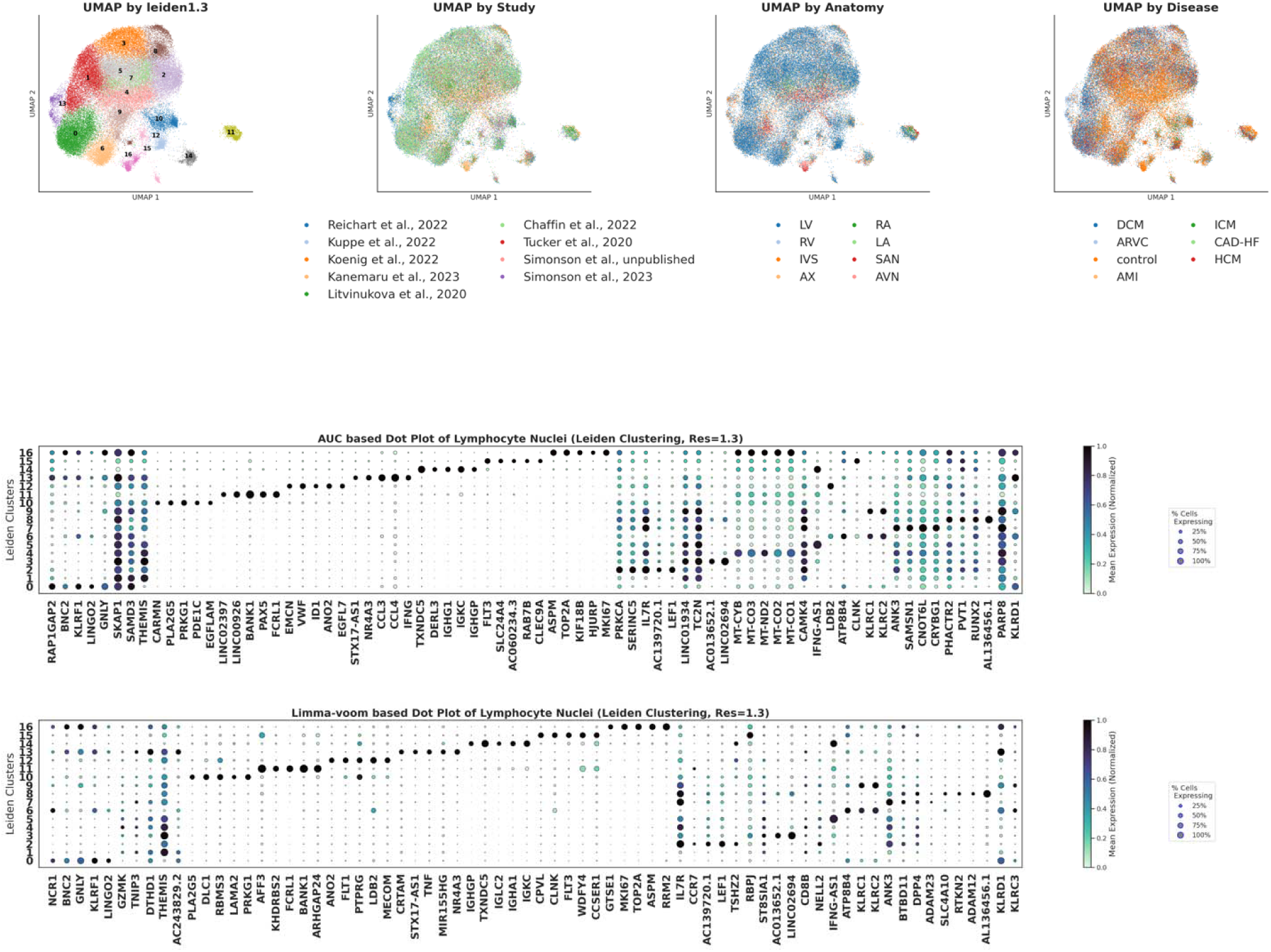
UMAP and Dot plot for HeartMap Lymphocyte nuclei. UMAPs are depicting Leiden-clustering resolutions, study, anatomy, and disease, where each point represents a nucleus, and colors differentiate the co-variate. Dot plots show the top AUC-score based and limma-voom based markers by log fold change for each Leiden cluster (see Methods).

**Supplemental Figure 14.**
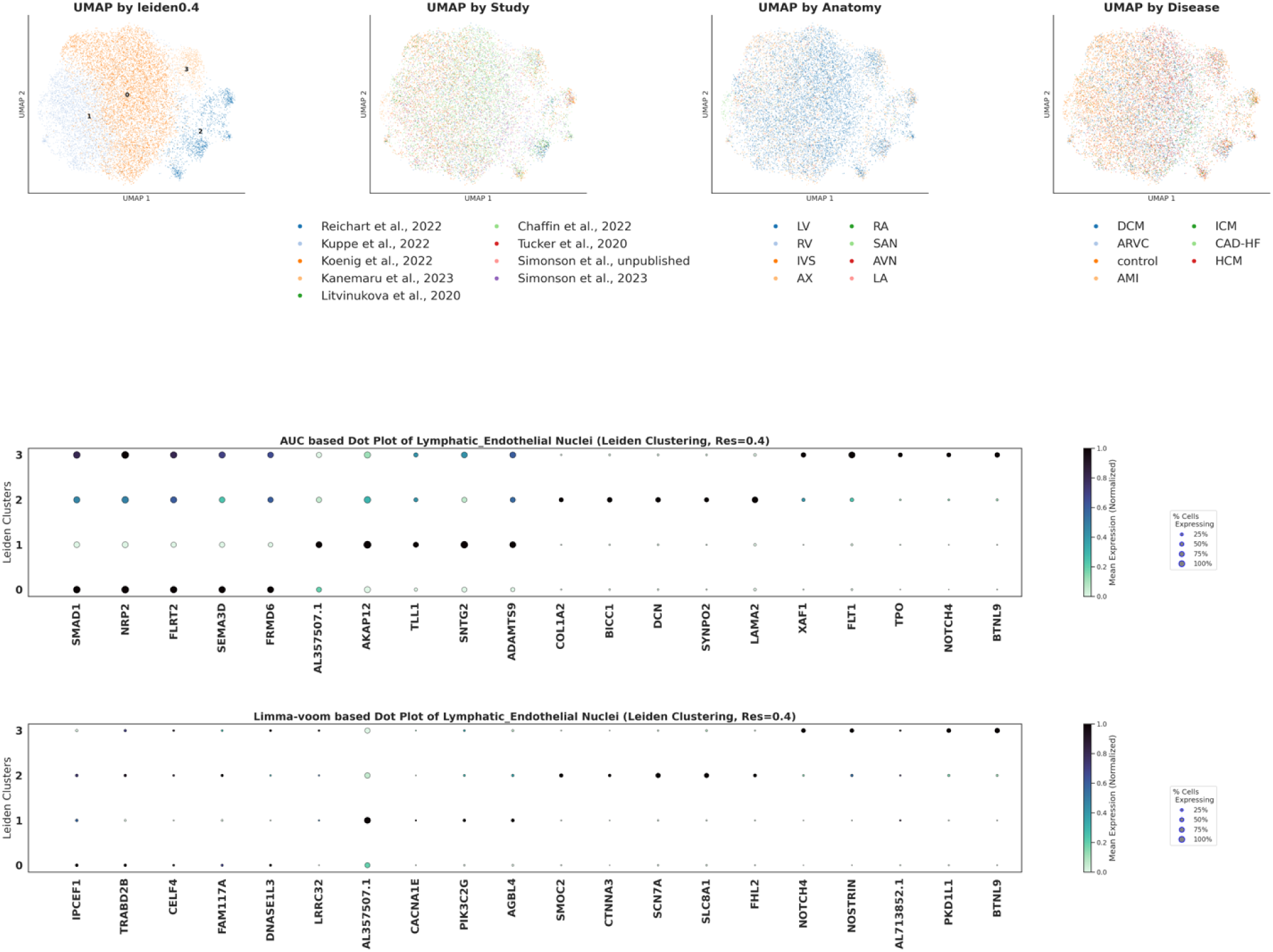
UMAP and Dot plot for HeartMap Lymphatic Endothelial nuclei. UMAPs are depicting Leiden-clustering resolutions, study, anatomy, and disease, where each point represents a nucleus, and colors differentiate the co-variate. Dot plots show the top AUC-score based and limma-voom based markers by log fold change for each Leiden cluster (see Methods).

**Supplemental Figure 15.**
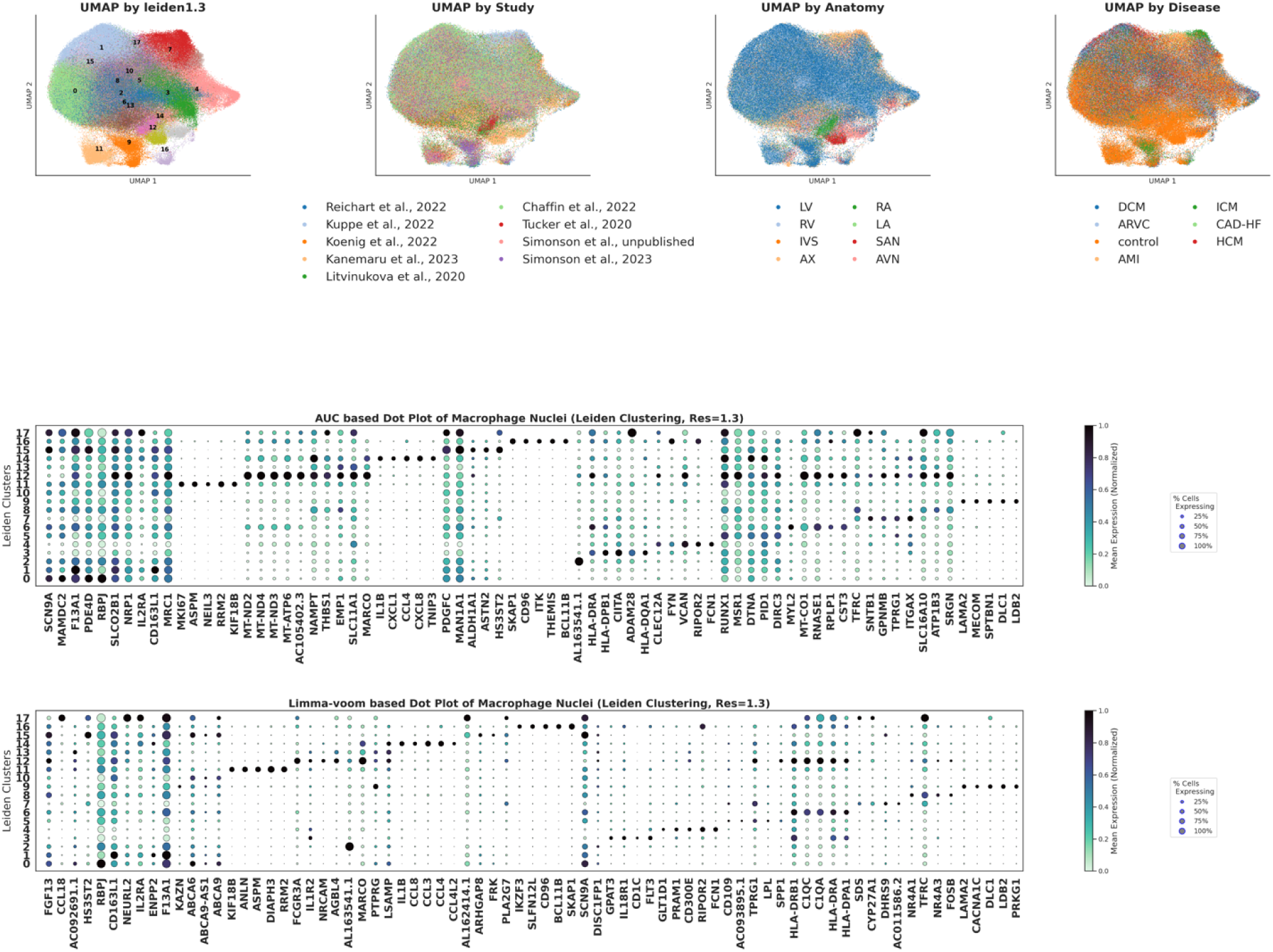
UMAP and Dot plot for HeartMap Macrophage nuclei. UMAPs are depicting Leiden-clustering resolutions, study, anatomy, and disease, where each point represents a nucleus, and colors differentiate the co-variate. Dot plots show the top AUC-score based and limma-voom based markers by log fold change for each Leiden cluster (see Methods).

**Supplemental Figure 16.**
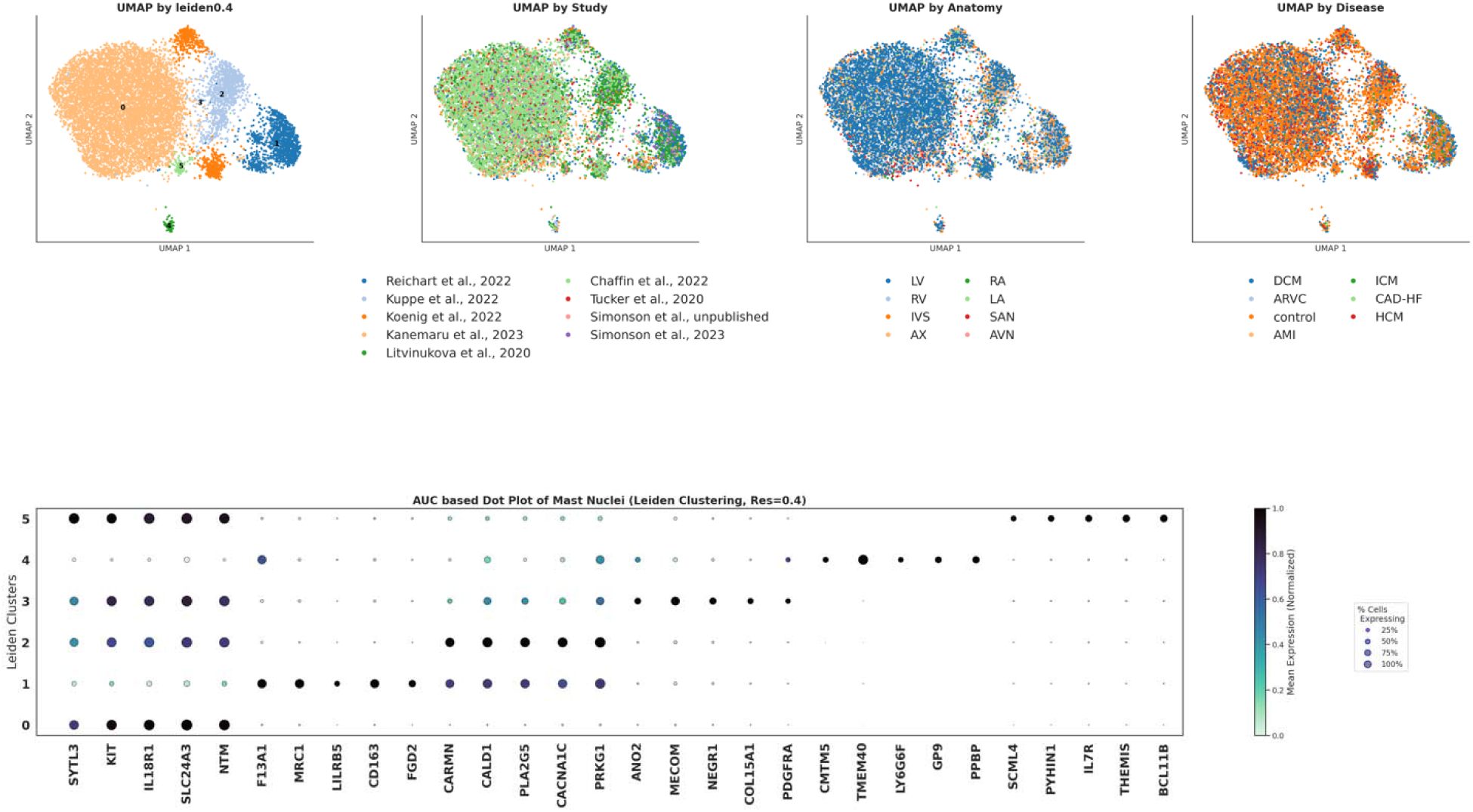
UMAP and Dot plot for HeartMap Mast Cell nuclei. UMAPs are depicting Leiden-clustering resolutions, study, anatomy, and disease, where each point represents a nucleus, and colors differentiate the co-variate. Dot plots show the top AUC-score based and limma-voom based markers by log fold change for each Leiden cluster (see Methods).

**Supplemental Figure 17.**
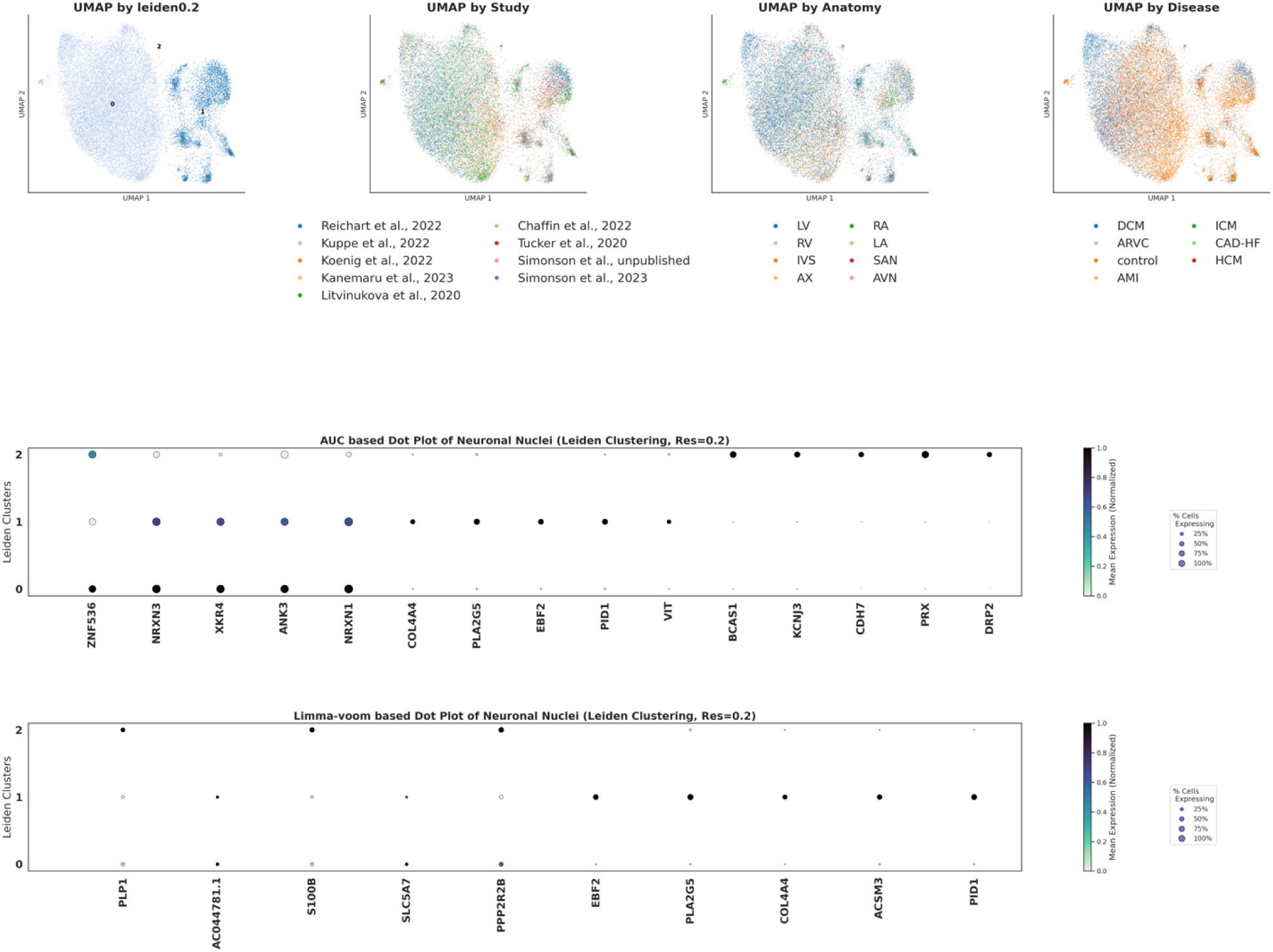
UMAP and Dot plot for HeartMap Neuronal nuclei. UMAPs are depicting Leiden-clustering resolutions, study, anatomy, and disease, where each point represents a nucleus, and colors differentiate the co-variate. Dot plots show the top AUC-score based and limma-voom based markers by log fold change for each Leiden cluster (see Methods).

**Supplemental Figure 18.**
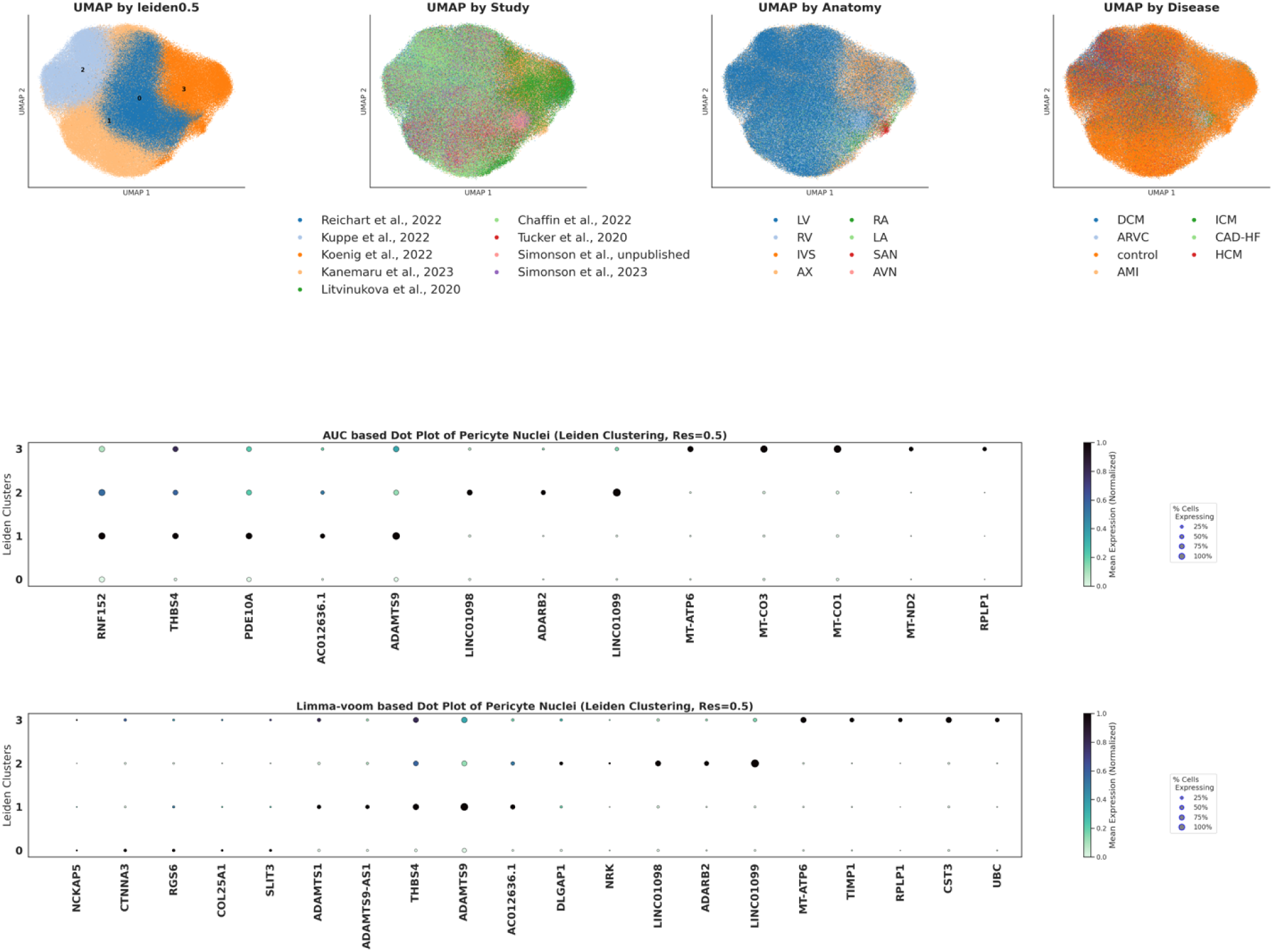
UMAP and Dot plot for HeartMap Pericyte nuclei. UMAPs are depicting Leiden-clustering resolutions, study, anatomy, and disease, where each point represents a nucleus, and colors differentiate the co-variate. Dot plots show the top AUC-score based and limma-voom based markers by log fold change for each Leiden cluster (see Methods).

**Supplemental Figure 19.**
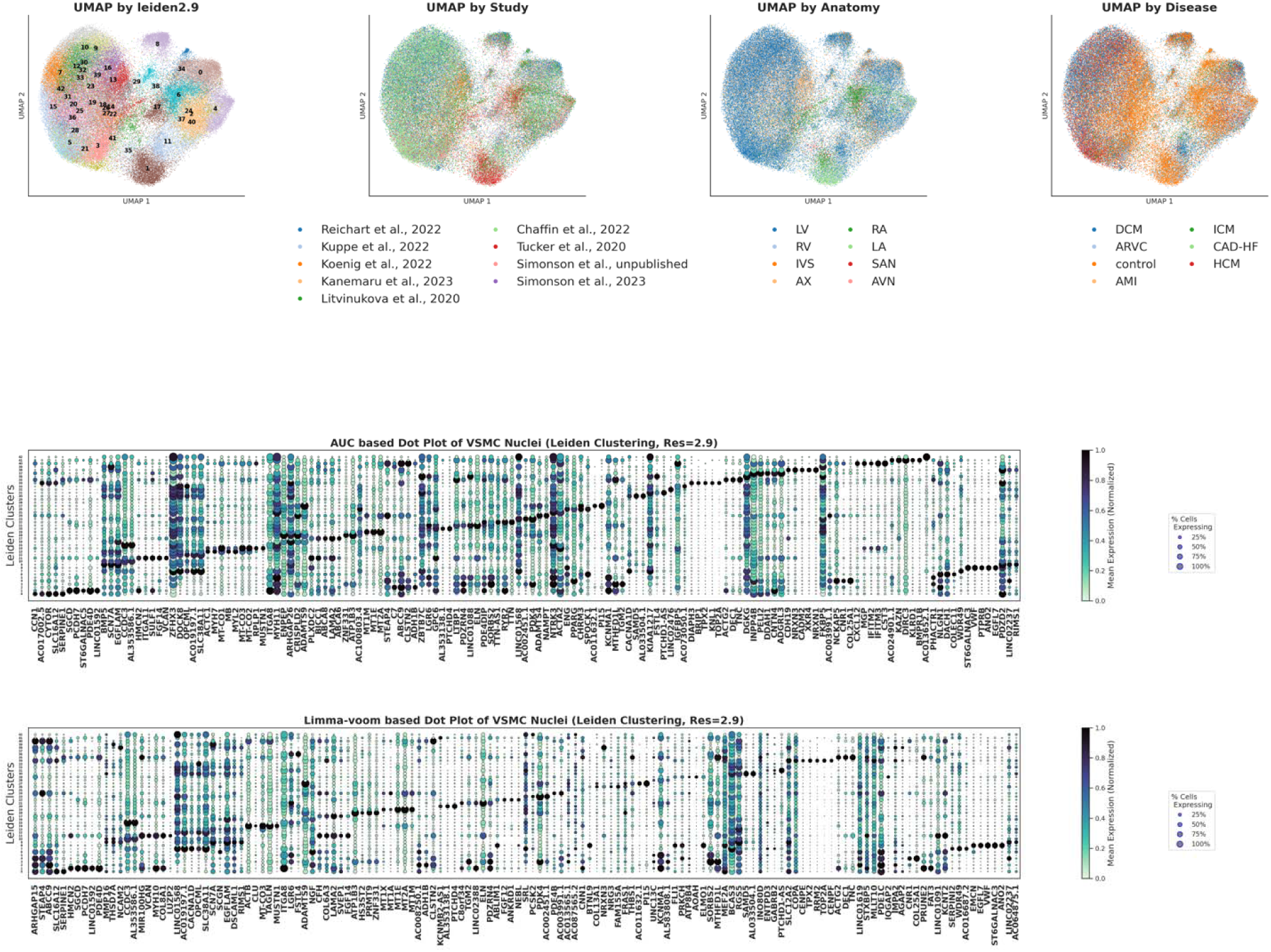
UMAP and Dot plot for HeartMap VSMC nuclei. UMAPs are depicting Leiden-clustering resolutions, study, anatomy, and disease, where each point represents a nucleus, and colors differentiate the co-variate. Dot plots show the top AUC-score based and limma-voom based markers by log fold change for each Leiden cluster (see Methods).

**Supplemental Figure 20.**
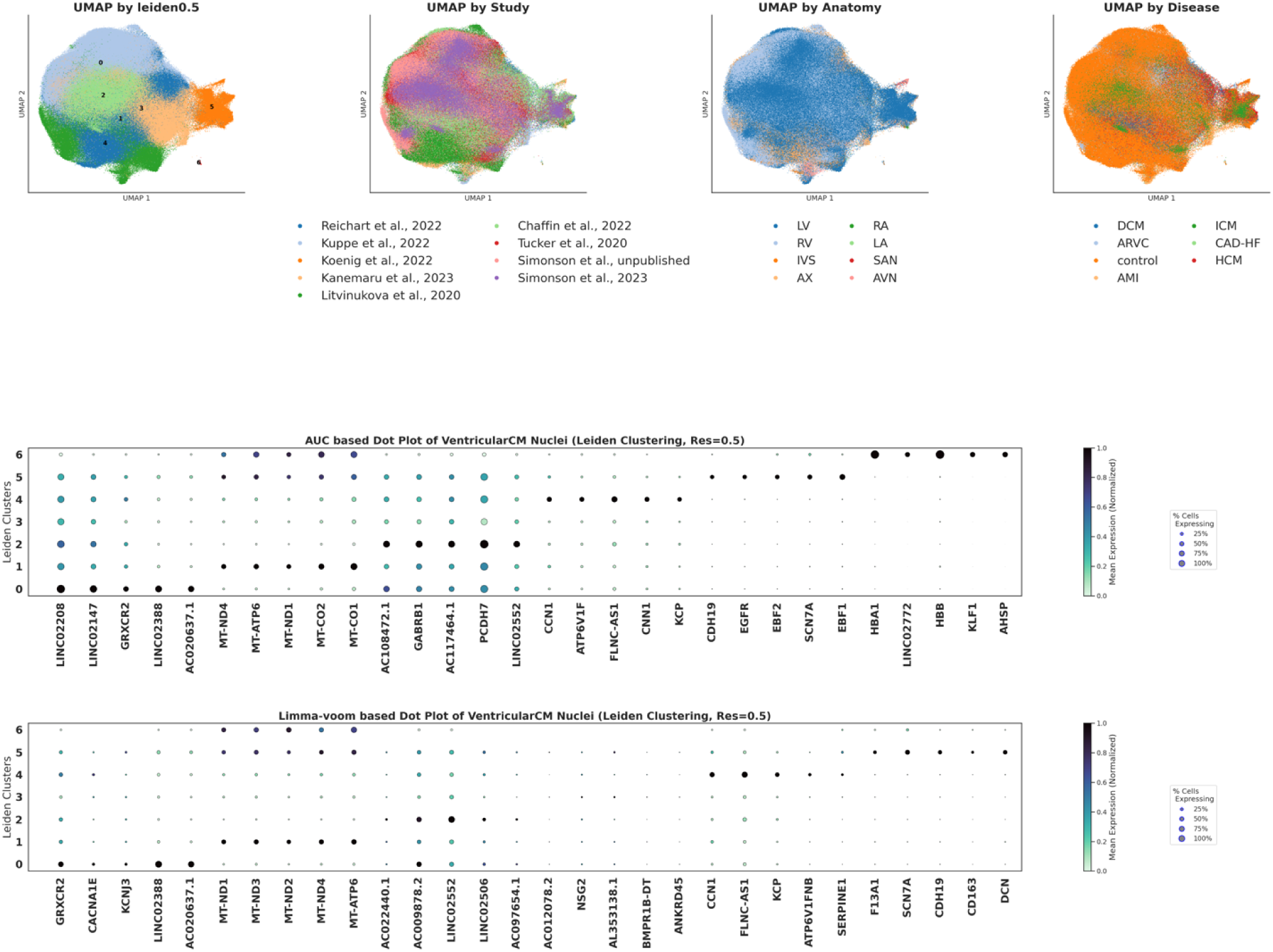
UMAP and Dot plot for HeartMap Ventricular Cardiomyocyte nuclei. UMAPs are depicting Leiden-clustering resolutions, study, anatomy, and disease, where each point represents a nucleus, and colors differentiate the co-variate. Dot plots show the top AUC-score based and limma-voom based markers by log fold change for each Leiden cluster (see Methods).

**Supplemental Figure 21.**
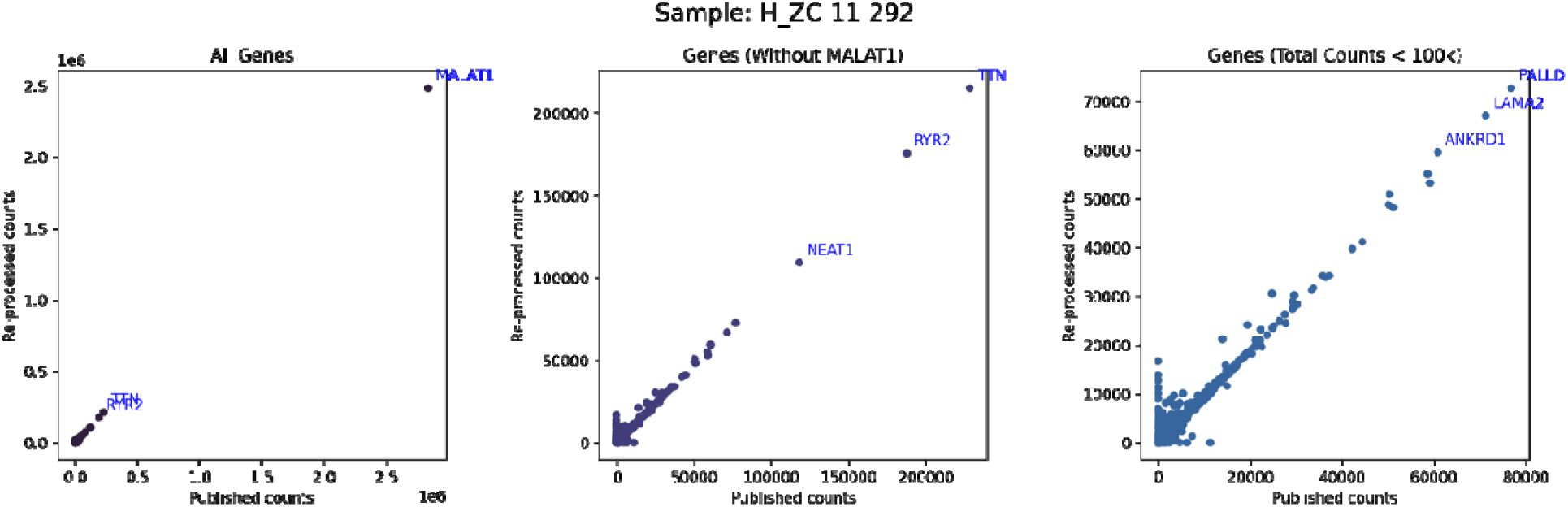
Example of correlation between re-processed and published gene counts for Koenig et al., 2022 sample. Correlation plots for the H_ZC-11-292 sample where each gene is plotted by counts in the re-processed and published studies demonstrating strong correlation. Only the shared 36,601 genes from the GRCh38 v.2020-A transcriptome are included.

## Supplemental Tables

**Supplemental Table 1. Marker genes for each of the cell types on the global map.** These were computed using the AUC-score based method (**see Methods**). Protein coding genes with an AUC > 0.6 are shown.

**Supplemental Table 2. Pairwise phenotypic comparisons between sex-assignments.** Genes are listed in each cell type with logFC, p.val. < 0.05, and the associated chromosome on which the gene is found.

**Supplemental Table 3. Shared genes between re-processed studies in the DCM vs NF controls analysis.** Strongly upregulated (logFC > 1.5) or downregulated (logFC < - 1.5) genes in the comparison of DCM to NF control across 3 studies. Only genes with a consistent effect direction across studies are shared. DCM, dilated cardiomyopathy; NF, non-failing; logFC, log_2_(fold-change); Background heuristic, a heuristic measurement of how likely the gene expression for the gene is likely driven by background RNA from other cell types; CM, cardiomyocyte; VSMC, vascular smooth muscle cell.

**Supplemental Table 4. Pathways enriched among the common genes in re-processed studies in the DCM vs NF controls analyses.** These pathways were computed using gene set enrichment (**see Methods**) with those listed having p. value < 0.05 across all enrichment scores per cell type.

**Supplemental Table 5. Gene set enrichment and ontology analysis for Fibroblasts.** Top 5 pathways by enrichment scores computed using gene set enrichment and gene ontology methods (**see Methods**) with those listed having p. value < 0.05.

**Supplemental Table 6. Percentage of activated fibroblast populations in each experimentally validated sample.** Computationally predictions and experimental findings for each of the nine samples that were in the image validation. Details on quantification calculation can be found in the Methods. NF, non-failing control; ICM, ischemic cardiomyopathy; HCM, hypertrophic cardiomyopathy; DCM, dilated cardiomyopathy.

**Supplemental Table 7.**
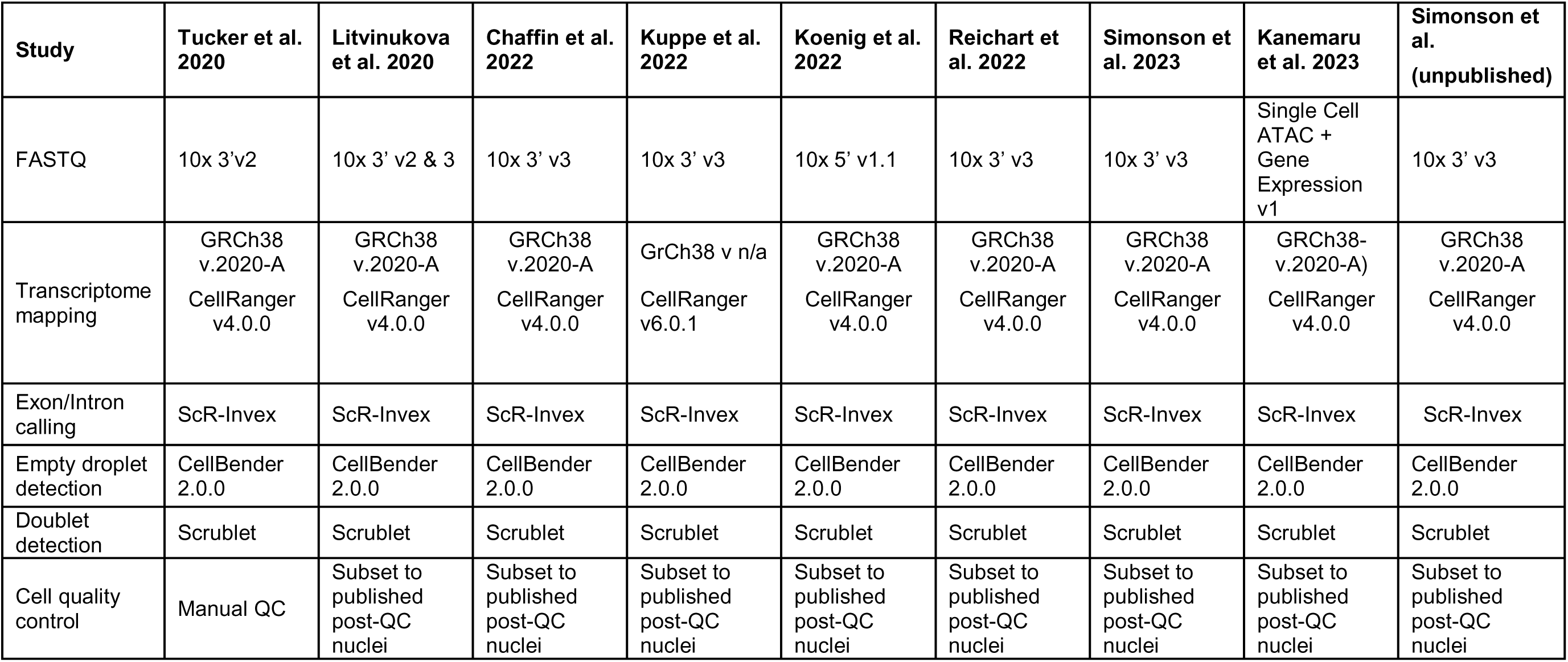
Pre-processing data and software packages used across all included data.

